# Iterative Bleaching Extends Multiplexity (IBEX) imaging facilitates simultaneous identification of all cell types in the vertebrate retina

**DOI:** 10.1101/2024.02.28.582563

**Authors:** Aanandita Kothurkar, Gregory S. Patient, Nicole C. L. Noel, Aleksandra M Krzywańska, Brittany J. Carr, Colin J. Chu, Ryan B. MacDonald

**Affiliations:** Institute of Ophthalmology, University College London, London EC1V 9EL, UK; Department of Ophthalmology and Visual Sciences, University of Alberta, Edmonton, AB, Canada T5H 3V9

**Keywords:** immunohistochemistry, zebrafish, development, neurons, glia

## Abstract

The vertebrate retina is a complex multicellular tissue made up of distinct neuron types and glia, arranged in a stereotypic layered organisation to facilitate vision. Understanding how these cell types come together to form precise circuits during development requires the ability to simultaneously discriminate between multiple cell types and their spatial position in the same tissue. Currently, we have a limited capacity to resolve all constitutive cell types and their relationships to one another due to our limited ability to combine multiple cellular markers. To extend this capacity, we have adapted a highly multiplexed immunohistochemistry technique known as Iterative Bleaching Extends Multiplexity (IBEX) and applied it to the development of the zebrafish (*Danio rerio)* retina. IBEX allows for multiple rounds of cellular labelling to be performed, before imaging and integration of data, resulting in the ability to visualise multiple markers on the same tissue. We have optimised IBEX in zebrafish using fluorescent micro-conjugation of known antibody markers to label the complete retina with up to 11 cell-specific antibodies. We have further adapted the IBEX technique to be compatible with fluorescent transgenic reporter lines, *in situ* hybridisation chain reaction (HCR), and wholemount immunohistochemistry (WMIHC). We then took advantage of IBEX to explore the multicellular relationships in the developing retina between glial cells and neurons and photoreceptor subtypes. Finally, we tested IBEX on retinas from the emerging ageing model, the killifish (*Nothobranchius furzeri)*, and developmental model, the African clawed frog (*Xenopus laevis),* demonstrating the usefulness of the technique across multiple species. The techniques described here can be applied to any tissue in any organism where antibodies are readily available to efficiently explore cellular relationships in the context of development, ageing or disease.

## INTRODUCTION

Tissues are made up of multiple cell types with regional and cell-specific molecular differences. To best understanding the relationships between these cells, and how they are altered during development, ageing and/or disease, we must have the ability to visualise the complete cellular landscape of a tissue. Our ability to do so relies on the accessibility of techniques to assay or discriminate between multiple cell types at the same timepoint using their transcriptomes, epigenomes, and/or proteomes as features to define cell states or interactions across an entire tissue (Choi *et al*., 2023). However, techniques used to study tissue composition and cellular relationships remain in refinement and can be costly to adapt for individual tissues or species of choice. Modern techniques, such as single-cell RNA-sequencing, can identify cell-specific molecular changes in individual cells on a large scale – however, the spatial organisation of these cells is lost in processing and must be mapped back onto the tissue to maximise their value. In contrast, immunohistochemistry (IHC) techniques allow visualisation of cellular proteins expressed in specific cell types in their undisturbed locations and thus provide spatial information. The number of antibody markers and, by consequence, the amount of information that can be obtained from a single tissue sample is limited by the number of fluorophores that can be imaged at one time. This is particularly relevant where few antibodies are validated and many are raised in the same host (e.g. rabbit), and so cannot be detected at the same time. Therefore, developing techniques to map the expression of multiple cellular markers onto a tissue will enhance our ability to understand cellular state, behaviour, and function in any tissue.

The retina is the light-detecting tissue at the back of the eye, comprised of several different types of neurons and glia. Retinal structure has been well-characterised since its early description by anatomists such as Cajal, who used dye labels to identify individual cell types and their organisation based on their unique locations and morphologies (Cajal, 1893). The highly organised retina is made up of five main neuronal cell types and a principal glia cell type called Müller glia (MG) (Fig. 1). Photoreceptors are the light-sensitive cells that synapse onto interneurons (horizontal cells, bipolar cells, and amacrine cells) which in turn, relay and modulate the signal and synapse with output neurons (retinal ganglion cells) that connect the retina to visual centres in the brain via the optic nerve. These cells are organised into three discrete cell layers – outer nuclear layer (ONL), inner nuclear layer (INL), and ganglion cell layer (GCL) – separated by two synaptic neuropils (outer and inner plexiform layers) and form relatively simple circuits (Masland, 2012). The zebrafish retina has been a useful model to study development and disease as it has a conserved organisation and cellular composition with other vertebrates (Avanesov and Malicki, 2010). Furthermore, it is an ideal neural tissue to image *in vivo* as the zebrafish eye is transparent during embryogenesis, develops rapidly (functional by 5 days post fertilisation (dpf) (Patterson *et al*., 2013), and has a full complement of cell-specific fluorescent reporter lines to visualise every cell type in the tissue (reviewed in Malicki *et al*., 2016). Further, there are a large number of antibodies with neuronal and glial specificity to discriminate different cell types and visualise morphology in the vertebrate retina (such as Yazulla and Studholme, 2001). However, studies of retinal development or disease remain constrained by our limited ability to combine these antibodies to visualise multiple cell types and directly assay cellular relationships or states in the same tissue at one time. Instead, such complex spatial information must be gathered by performing IHCs with numerous combinations of the same antibody pools across many tissue sections.

**Figure 1.**
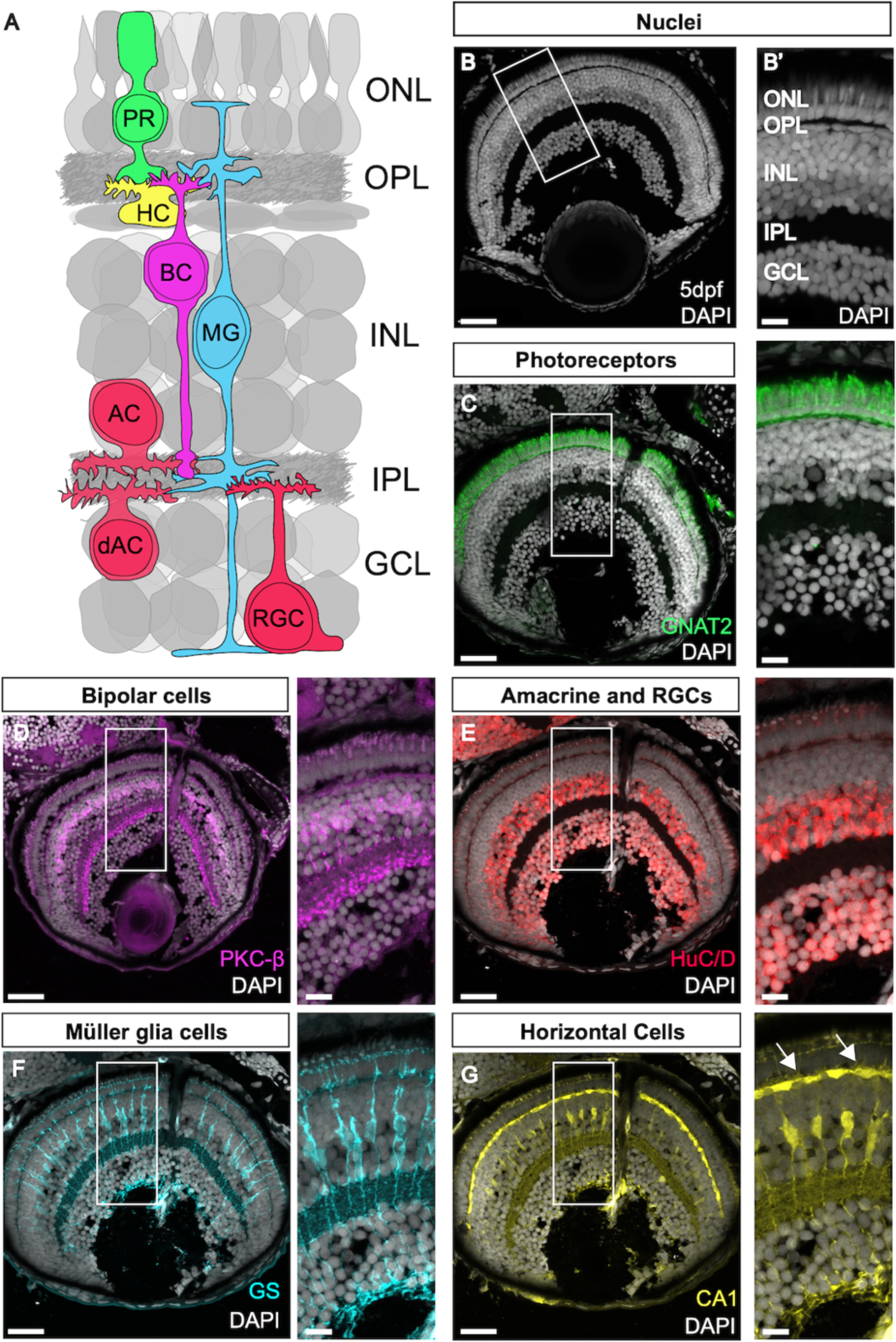
The retina is made up of highly organised layers composed of neurons and glia. (A) Schematic of the retina showing the layers and major cell populations with each cell type colour coded. (B) DAPI staining showing the nuclear layers of the retina. (B’) Zoom of B. (C-G) Antibody staining for the main cell types in the zebrafish retina. Arrows in G show horizontal cells. OLM: outer limiting membrane, ONL: outer nuclear layer, OPL: outer plexiform layer, INL: inner nuclear layer, IPL: inner plexiform layer, GCL: ganglion cell layer. Scale bars - 25µm for whole retina, 10µm for zoom images.

Here, we overcome these challenges by adapting and expanding the Iterative Bleaching Extends Multiplexity (IBEX) technique in the vertebrate retina (Fig. 2). IBEX is a technique developed in mouse and human tissues that allows simultaneous visualisation of up to 60 markers on a single tissue sample (Radtke *et al*., 2020), thereby providing large-scale, detailed multicellular spatial analysis of tissue. It relies on fluorescently conjugated primary antibodies to enable use of multiple antibodies raised in the same species while avoiding cross-reactivity and permits bleaching of signal to conduct sequential rounds of immunolabelling. First, we validated “micro-conjugations” whereby a small volume of antibody is directly linked to fluorescent dyes to overcome the critical issue of multiple antibodies raised in the same species. Importantly, these fluorophores can be bleached and are compatible with multiple rounds of IHC required for the IBEX technique. Using IBEX, we then labelled every cell type in the retina with 11 specific antibody markers. We enhanced the capabilities of the IBEX technique in zebrafish by pairing with cell-specific transgenic reporter lines, wholemount IHC (WMIHC), and *in situ* hybridisation chain reaction (HCR) in zebrafish. We also use IBEX to describe the development of two key cell types in the retina: photoreceptors and MG. The techniques described here will be valuable for any tissue and are applicable to any other study where multiplexed IHC is required. Finally, to demonstrate the usefulness of IBEX across species we applied it to two additional aquatic vertebrates: the African clawed frog (*Xenopus laevis)* and the African turquoise killifish (*Nothobranchius furzeri)*. Both models are currently constrained by the lack of tools, such as few established transgenic reporter lines, making the ability to visualise multiple-cell types in the retina challenging. As such, we have adapted the IBEX technique to expand our ability to visualise multiple cell types in the same tissue sample to explore cellular relationships in the developing vertebrate retina.

**Figure 2.**
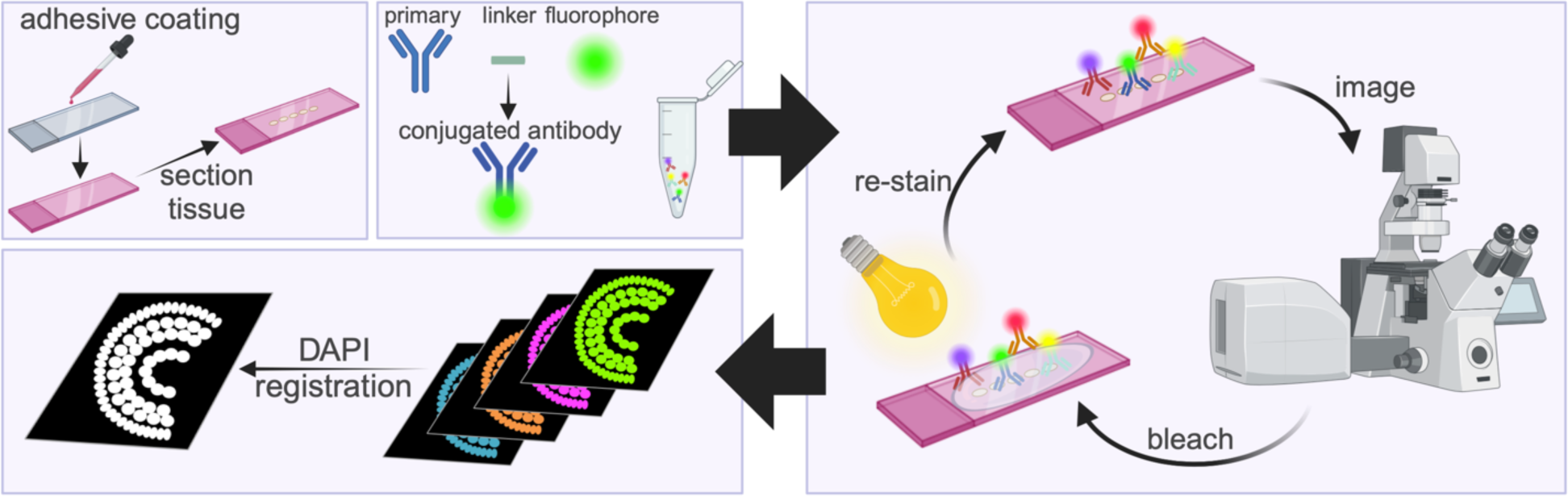
Schematic of the IBEX method. Slides are coated with chrome alum gelatin to prevent tissue lost, then tissue sectioned onto slides. Antibodies are micro-conjugated by mixing the primary antibody with a linker and fluorophore. The antibodies are applied to the slide, incubated, imaged, then bleached using bright light and lithium borohydride before being re-stained. After the imaging rounds are completed, nuclear stains (DAPI) are used to register the image, allowing for all stains to be visualised together. Made with Biorender.

## RESULTS

### Direct conjugation to fluorophores facilitates labelling with multiple antibodies raised in the same host on the same tissue

A major hurdle in IHC is labelling with multiple antibodies raised in the same animal (e.g. rabbit), as it would not be possible to distinguish between the antibodies using traditional secondary antibodies. To overcome this limitation, we used “micro-conjugation” reactions (see Methods) to directly link primary antibodies with distinct fluorophores, avoiding use of secondary antibodies and gaining the flexibility to label each antibody with a fluorophore of choice (Fig. S1). To ensure there was no cross-reactivity or quenching of signal due to competitive antibody binding, we conjugated the rabbit GNAT2 antibody, which labels cone photoreceptors in the zebrafish retina, with four different fluorophores. Using confocal microscopy, we observed robust signal for each of the fluorophores in the photoreceptor layer with no noticeable loss of signal due to multiple conjugated antibodies against the same protein (Fig. S1).

It is critical for the multiplexity of the IBEX technique to be able to quench the fluorescent signals between rounds of IHC. We demonstrate that CoraLite fluorophores can be successfully bleached using lithium borohydride (LiBH_4_) and show a near complete loss of signal and no autofluorescence in any channel post bleaching (Fig. S1). Next, to determine if we could concurrently label cells with four distinct rabbit polyclonal antibodies, we micro-conjugated each with different fluorophores and conducted a single round of IHC and imaging (Fig. 3). We labelled cone photoreceptors with GNAT2, bipolar cell ribbon synapses with Ribeye-A, bipolar cell terminals with PKC-β, and MG with RLBP1; we were able to visualise these markers concurrently without any noticeable cross-reactivity. Therefore, using micro-conjugations we can reliably visualise and quench the fluorophores of antibodies raised in the same species on the same tissue section.

**Figure 3.**
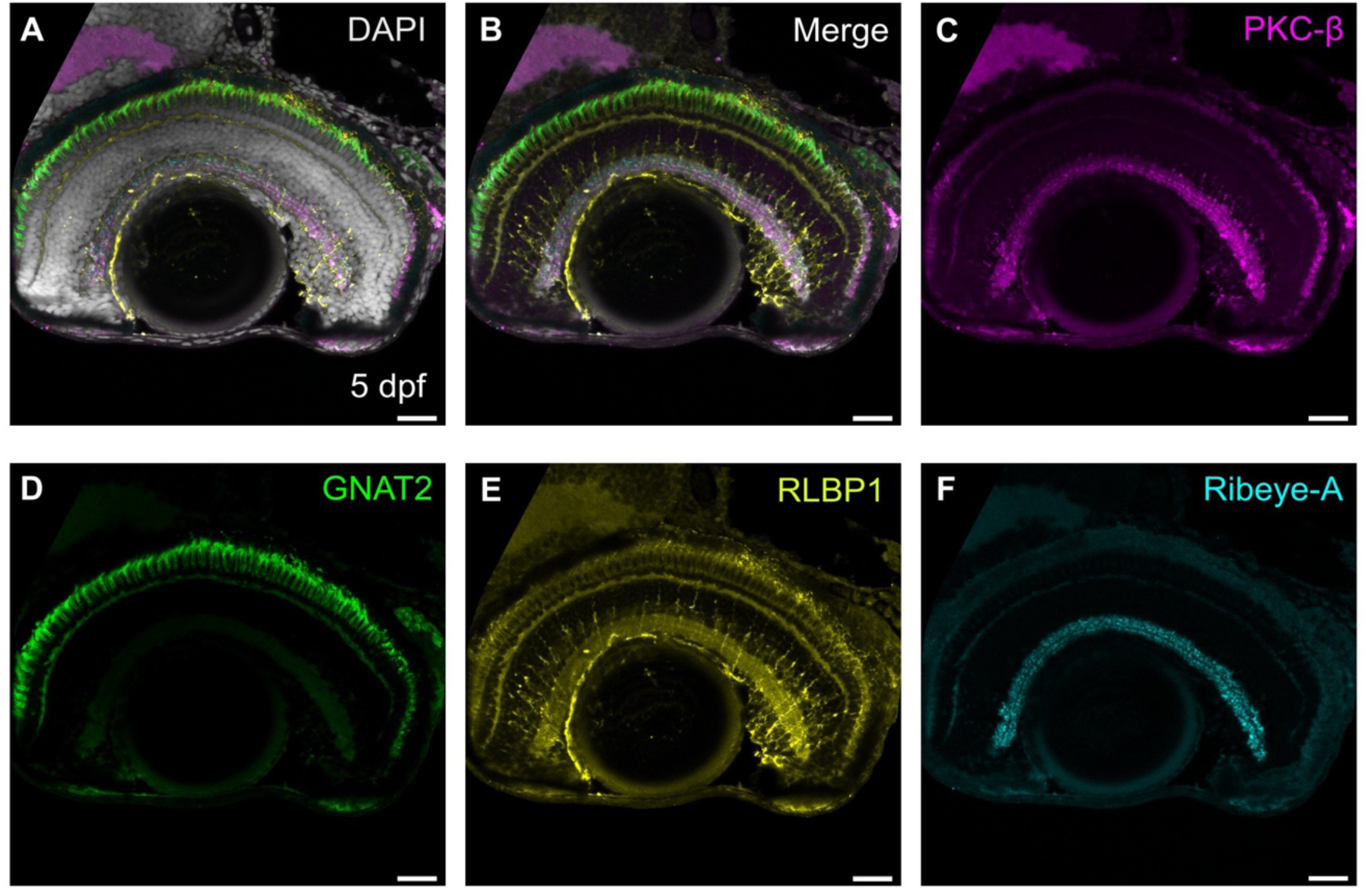
Direct conjugation to fluorophores facilitates multiple single species antibody labels on the same tissue. (A) Merged epifluorescence images of a single sagittal retinal section immunolabelled with four antibodies raised in rabbit: PKC-β (magenta), GNAT2 (green), RLBP1 (yellow), Ribeye-A (cyan), and nuclear stain DAPI (grey). (B) Merged image without DAPI. (C) Retinal section immunolabelled with PKC-β, marking bipolar cells. (D) Retinal section immunolabelled with GNAT2 marking cones. (E) Retinal section immunolabelled with RLBP1, marking Müller glia cells. (F) Retinal section immunolabelled with Ribeye-A marking ribbon synapses. Scale bars - 25µm

### Adapting IBEX to label all cell types in the zebrafish retina

To label every major cell type in the retina with IHC, we designed a panel of markers against proteins expressed in each cell type of the zebrafish retina composed of micro-conjugated antibodies and directly conjugated antibodies. First, we optimised each antibody for use with micro-conjugation by testing for bright, specific labelling in single IHC tissue staining and bleaching (see Materials and Methods). In some cases, directly conjugated antibodies did not show sufficient staining to be easily visualised by standard IHC or they were not successfully bleached by LiBH_4_. Fortuitously, a standard antigen retrieval step was sufficient to bleach and enhance the staining for these antibodies. Using three rounds of iterative bleaching followed by standard confocal imaging, we labelled each cell type in the retina using 11 markers and DAPI in the same tissue section (Fig. 4, Supplementary Video 1). In each IHC round, we used DAPI to label nuclei, which is used for alignment and the ultimate integration of multiple markers on the same tissue as it does not bleach. This provides a consistent fiducial landmark for image registration. The open source SimpleITK registration software (Radtke *et al.,* 2020) for registration of confocal Z-stacks is effective at increasing the alignment of DAPI signal between the three rounds (Fig. S2) and allows channels from the different rounds to be merged. As such, we developed panels of combinatorial fluorescent antibody labels against each cell type in the retina, imaged each panel in successive imaging rounds after quenching of fluorophores, and integrated the data onto a single image file (Fig. 4A). To increase the rate at which we can acquire data from multiple samples, we also optimised IBEX and the panel of markers for an epifluorescence imaging system with onboard deconvolution. This allows for a large area of tissue to be imaged quickly and effectively over multiple rounds to visualise 9 antibodies and a lectin stain on the same tissue (Fig. S3).

**Figure 4.**
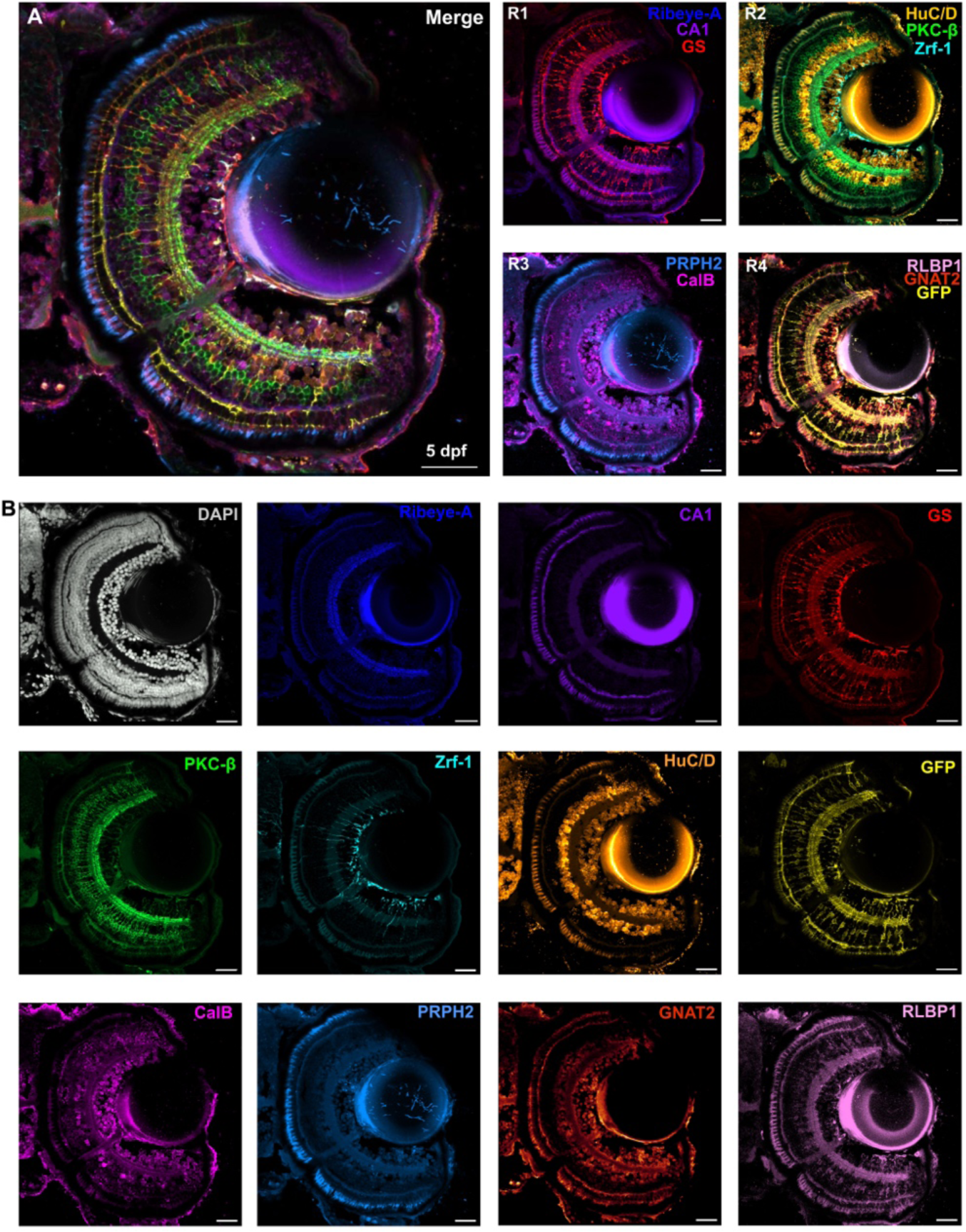
IBEX enables simultaneous labelling of all retinal cell types. (A) Confocal images of 5 dpf Tg(*tp1:eGFP:CAAX*) zebrafish retina showing different rounds of immunolabelling (R1-4) using IBEX and the merge composite image of each of these rounds. R1 was carried out using Alexa Fluor secondary antibodies, while other rounds used microconjugated antibodies. (B) Confocal images showing each antibody used in (A) to immunolabel a single sagittal retinal section with DAPI and 11 different markers: Ribeye-A (dark blue), carbonic anhydrase (CA1, purple), glutamine synthetase (GS, red), PKC-β (green), Zrf-1 (cyan), HuC/D (orange), GFP transgene (yellow), calbindin (CalB, magenta), peripherin-2 (PRPH2, light blue), GNAT2 (orange), and RLBP1 (pink). Scale bars - 25µm.

### IBEX is compatible with cell-specific transgenic reporter lines

We aimed to assess whether we could combine IBEX with existing zebrafish transgenic lines to enhance our multiplex toolbox. To incorporate transgenic reporter lines into the IBEX technique, it is ideal for the fluorescent protein to bleach and make the channel available for future imaging rounds. We tested whether endogenous fluorescent protein signals could be quenched and then re-labelled with fluorescent protein-specific antibodies by IHC. We tested several transgenic lines containing cytosolic (Tg(*GFAP:GFP*),Tg(*vsx1:GFP*)^nns5^, Tg(*TP1:Venus-Pest*),Tg(*ptf1a:dsRed*)^ia6^, Tg(*rho:YFP*)^gm500^) or membrane-targeted (Tg(*tp1bglob:eGFP-CAAX*)) fluorescent proteins. We found that both cytosolic and membrane-tagged GFP bleached after antigen retrieval methods (Fig. S4). These transgenes could then be boosted with an anti-GFP antibody and imaged in the last round of labelling. We confirmed that the transgene retained its cell specificity during this process by co-labelling and observing overlapping labelling with a cell specific antibody, PKC-β for Tg*(vsx1:GFP)* and GS for Tg*(tp1bglob:eGFP-CAAX)* (Fig.S4 A’’,A’’’, B’’,B’’’). However, we could not bleach the RFP or YFP transgenic lines with LiBH_4_ in combination with intense light nor sodium citrate antigen retrieval (Fig. S4C, D).

### IBEX can be combined with fluorescent *in situ* hybridisation

Antibodies specific for a cell type or protein of interest can be limited in zebrafish. As an alternative, *in situ* hybridisation chain reaction (HCR) is a robust method to label mRNA of interest in zebrafish (Choi *et al*., 2010, 2016, 2018). HCR has been previously combined with IHC (Howard et al., 2021; Ibarra-García-Padilla et al., 2021; Ćorić et al., 2023), however, these are limited by the number of channels available in a single labelling round on standard microscopes. With a view to overcoming this limitation, we tested whether *in situ* hybridization chain reaction (HCR) methods to label mRNA would be compatible with IBEX, such that we could conduct a multiplex HCR followed by bleaching, IHC and integration of labelling techniques on the same retina. For this, we performed HCR for three genes of interest: *cyp26a1*, *glula*, and *vsx1.* The expression of these genes is known to be specific to different retinal cell populations: MG (*cyp26a1* and *glula*) and bipolar cells (*vsx1*) (Fig. 5A-A′′′′). We then attempted to quench the signal of these fluorophores using LiBH_4_ treatment. However, we did not observe a significant reduction in signal for Alexa Fluor 555 (Fig. 5B-B′′′) We were able to bleach this signal using the antigen retrieval technique (Fig. 5B′′′′) before conducting a subsequent round of IHC with MG and bipolar cell antibody markers (Fig. C-C′′′′). We overlayed these two rounds of imaging, one HCR and one IHC, which allowed us to visualise expression of the three transcripts of interest and confirm co-localisation with different retinal cell populations labelled by antibodies (Fig. 5D-D′′′′).

**Figure 5.**
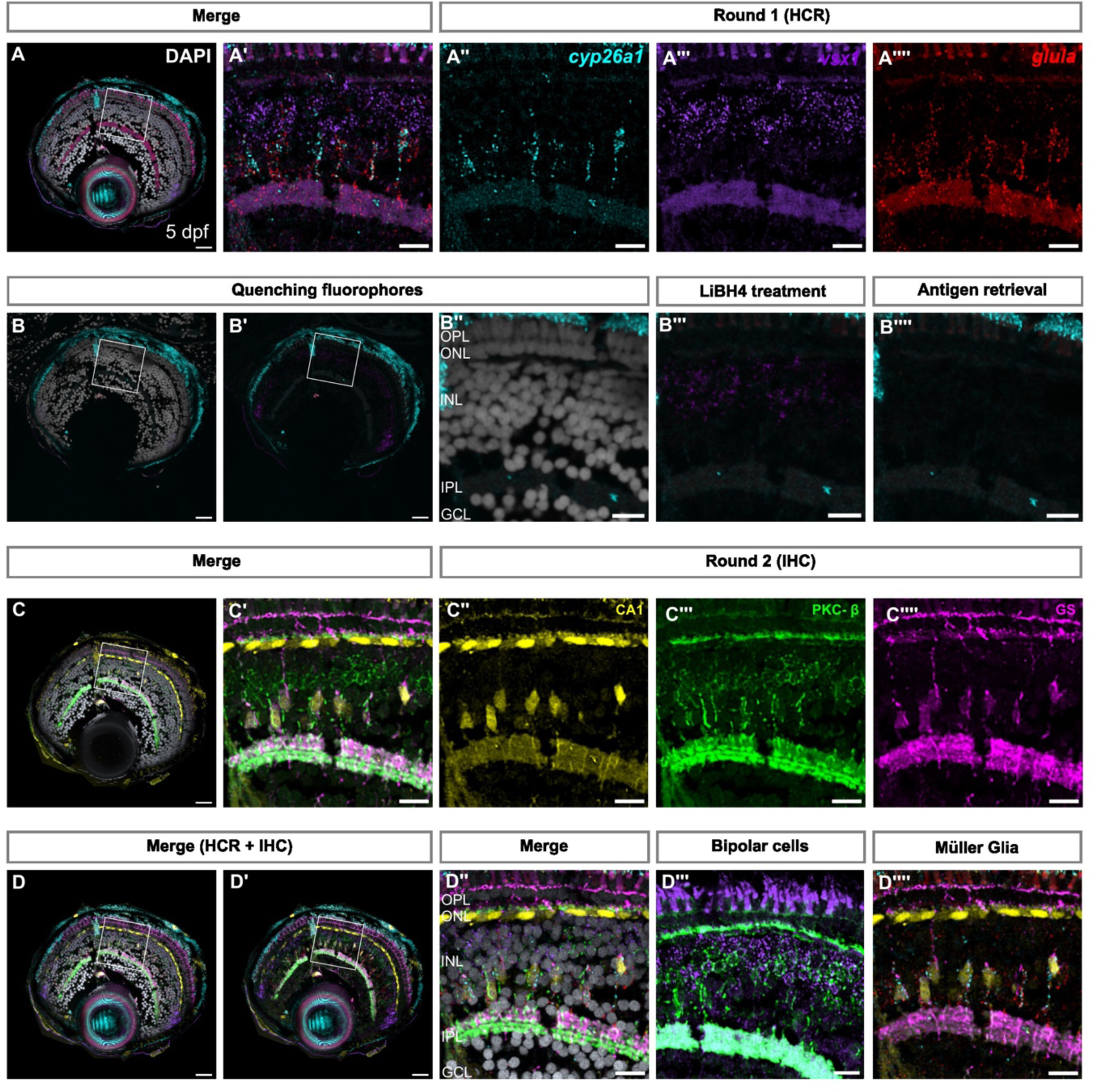
IBEX is compatible with fluorescent *in situ* hybridisation chain reaction. (A) Confocal images of retinal sections showing mRNA expression of *cyp26a1*, *glula*, and *vsx1*, using *in situ* hydridisation chain reaction (HCR). (A′-A′′′′) Zoom on the region of interest indicated in (A). (B-B′′′) Confocal images showing reduced signal of Alexa Fluor-488 and Alexa Fluor-647, but not Alexa Fluor-555 after LiBH4 treatment. (B′′′′) Heating in sodium citrate at 60°C causes inactivation of Alexa Fluor 555 as well as Alexa Fluor 488 and Alexa Fluor 647. (C) Confocal images of retinal sections immunolabelled with CA1 (yellow), PKC-β (green), and GS (magenta). (C′-C′′′) Zoom on region of interest indicated in (C). (D) SimpleITK registered image, showing overlay of both rounds of imaging, and overlay of *in situ* probes and antibodies detecting Müller glia and bipolar cells, respectively. (D′′-D′′′) Zoom of region of interest shown in (D,D′). GCL: ganglion cell layer, IPL: inner nuclear layer, ONL: outer nuclear layer, OPL: outer plexiform layer. Scale bars - 25µm for whole retina, 10µm for zoom images.

### Wholemount IBEX facilitates whole tissue labelling in zebrafish

The relatively small size of the zebrafish retina and the ability to treat the fish to make them optically transparent lends itself to WMIHC (Inoue and Wittbrodt, 2011; Santos, Monteiro and Luzio, 2018). This technique facilitates the study of cell structure and shape in its native conformation and overcomes the potential disruption of cell morphology and tissue damage introduced by cryosectioning. However, the hurdle of visualizing multiple cell types in the same sample remains. Therefore, we tested whether the micro-conjugated antibody staining and bleaching is compatible with thicker tissues in WMIHC before carrying out the IBEX protocol. We tested the protocol by immunolabelling MG and photoreceptors with two different antibodies each (GS and zrf-1 for MG, and GNAT2 and blue opsin for photoreceptors), over two successive rounds of imaging (Fig. 6A,A′,C,C′). Micro-conjugated antibodies penetrated the tissue and specifically labelled photoreceptors and MG. Treatment with LiBH_4_ successfully bleached the signal of each of the fluorophores between rounds (Fig. 6B). The SimpleTK registration software allowed us to combine the images and observe the co-localisation of antibodies labelling Müller glia and photoreceptors, respectively, across different rounds of immunofluorescence (Fig. 6E,E′,F, Supplementary video 2). Therefore, WMIHC when combined with IBEX allows 3D labelling of multiple cell types and alignment of their spatial relationships to one another between rounds.

**Figure 6.**
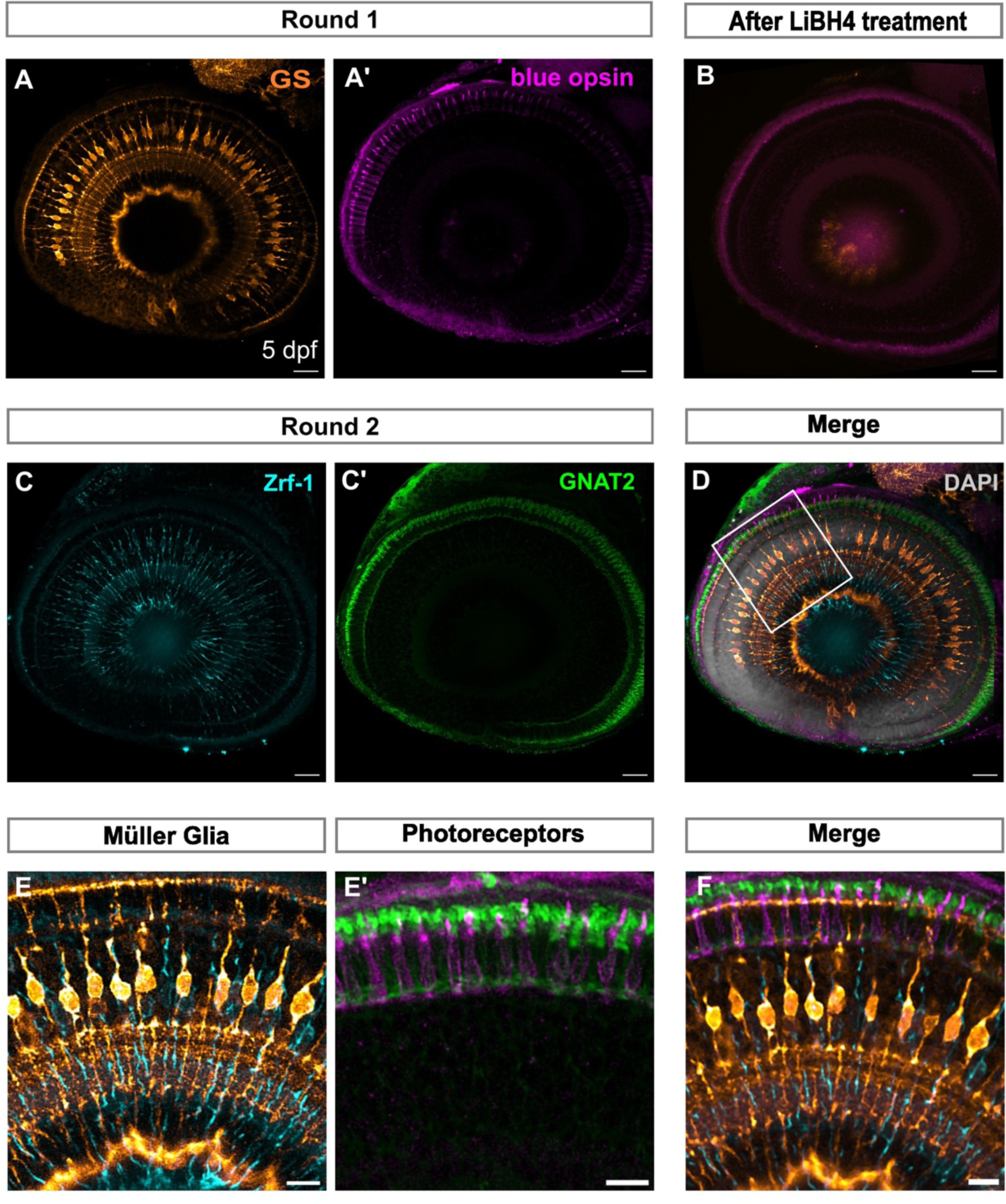
Wholemount IBEX facilitates whole tissue labelling in zebrafish. (A,A′) Confocal images of wholemount zebrafish larvae at 5 dpf immunolabelled with GS (orange) and blue opsin (magenta). (B) Tissue after bleaching with LiBH4 showing reduced signal of fluorophores CoraLite 488 and CoraLite 647. (C,C′) Confocal images of the second round of immunolabelling to detect Zrf-1 (cyan) and GNAT2 (green). (D) Merge of both rounds of immunolabelling using SimpleTK registration pipeline counterstained with DAPI (grey). (E) Overlap of MG markers GS and Zrf-1 across round 1 and 2. (E′) Overlap of photoreceptor markers blue opsin and GNAT2 across round 1 and 2. (F) Zoom in of (D), showing overlay of MG and photoreceptor labelling. dpf: days post fertilisation, MG: Müller glia. Scale bars - 25µm for whole retina, 10µm for zoom images.

### IBEX facilitates the characterisation of retinal histogenesis and patterning

The retina has a stereotyped histogenesis whereby retinal neurons and glia are born and specified in distinct temporal sequence during retinogenesis (Agathocleous and Harris, 2009). Specification of the zebrafish retina begins at 24 hours post fertilisation (hpf) as a retinal primordium, completing histogenesis by 73 hpf (Easter and Nicola, 1996) with robust vision beginning at 5 dpf. We used this well-characterised developmental pattern to determine the utility of IBEX to describe cellular morphologies in the highly dynamic developing retina. We focussed on two main cell types: photoreceptors, which have five distinct subtypes that are challenging to visualise simultaneously by traditional methods, and MG, due to their highly dynamic morphological changes across retinal development. We used cryosections at different key timepoints of retinal development to accomplish this.

Zebrafish photoreceptors undergo rapid development, with light-sensitive opsin mRNA expression detectable by 60 hpf (Robinson, Schmitt and Dowling, 1995). Zebrafish are tetrachromats: they have rods and four cone photoreceptor subtypes, maximally sensitive to ultraviolet (UV), blue, green, and red light. Different photoreceptor types are identifiable by specific markers; however, traditional methods make it challenging to label all photoreceptor subtypes such that they are distinguishable from one another. We combined antibody labelling with a transgenic line with fluorescently labelled rods (Tg(*rho:YFP*) line) to label all photoreceptors with subtype resolution in the developing zebrafish retina at three stages (3, 4, and 5 dpf) (Fig. 7). Of note, YFP did not bleach with LiBH4 nor antigen retrieval, and therefore antibody labelling rounds were adjusted to avoid use of 488 fluorophores. We distinguished between the cone subtypes by utilising antibodies against UV, blue, and red opsin (via 1D4), as well as arrestin 3a (with zpr-1). Zpr-1 labels both red and green cones; green cones can therefore be identified as cells that are arrestin 3a-positive but do not stain for red opsin (Fig. S6). At 3 dpf, developing cone photoreceptors stain with GNAT2 and zpr-1 (Fig. S5A′′), and have small outer segments labelled with antibodies for PRPH2, UV opsin, blue opsin, and red opsin (Fig. S5A′′′, A′′′′). Most of the cones with discernible outer segments were observed in the central retina. Few newly developed YFP-positive rods can also be observed in the retinal periphery. By 4 dpf, cones appear more morphologically mature with lengthened outer segments (Fig. S5B-B′′′′). As mentioned, zebrafish cones are functional by 5 dpf and the animals begin to perform complex visually mediated behaviours, such as prey capture (Patterson *et al*., 2013). Corresponding with this, 5 dpf zebrafish cones have visually longer outer segments compared to 4 dpf with a tapered morphology (Fig. S5C-C′′′′). Furthermore, there appear to be phagosomes in the RPE staining for zpr-1, GNAT2, UV opsin, and PRPH2, suggesting that there is outer segment disc shedding at this stage.

**Figure 7.**
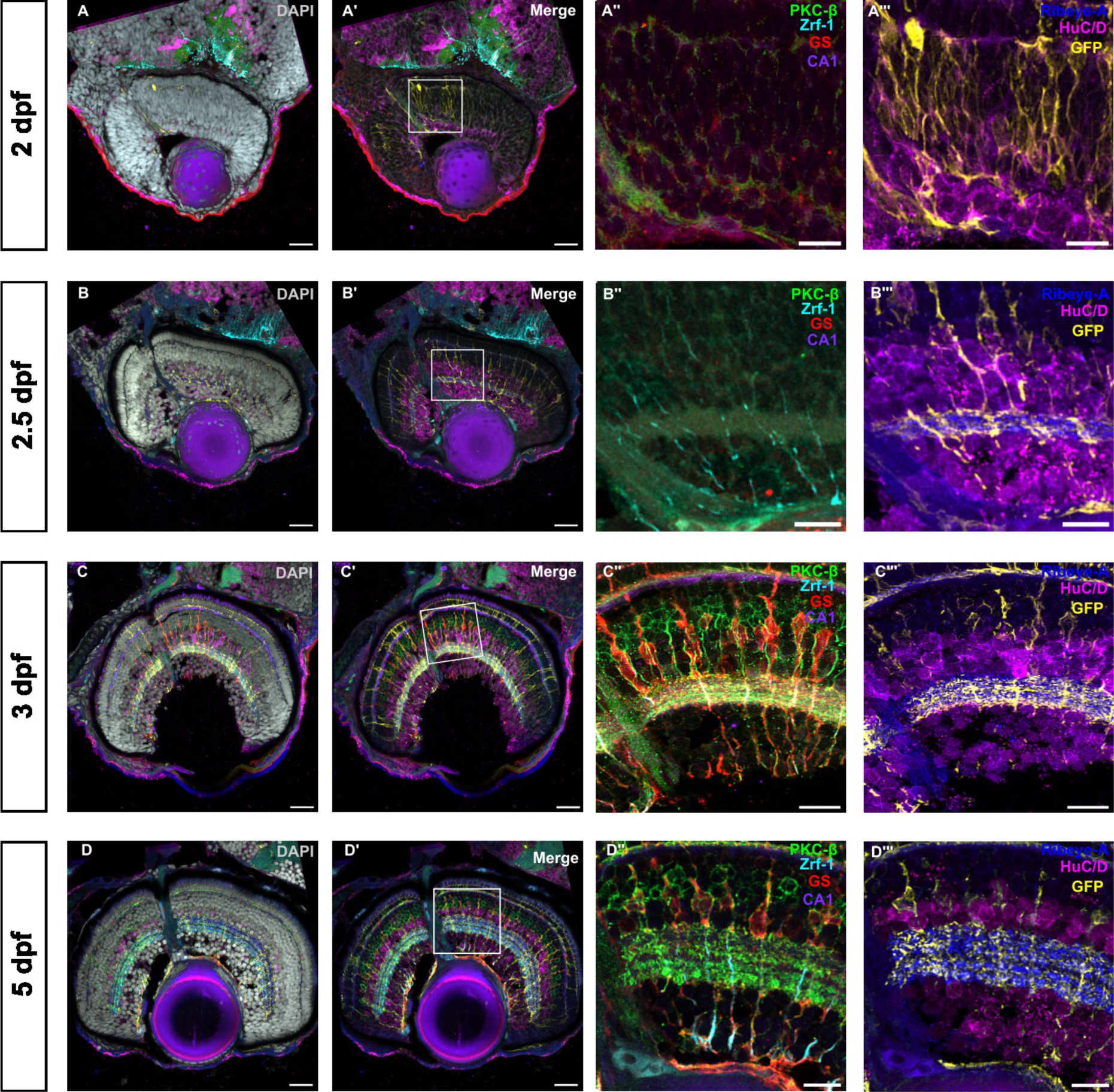
Visualisation of glial and neuronal development in the zebrafish retina. Confocal images of the developing zebrafish retina from 2 dpf to 5 dpf immunolabelled with DAPI and 7 different markers using IBEX over 3 rounds of immunolabelling with IBEX. A, B,C,D show merge of all 7 markers with nuclear stain DAPI, while A′,B′,C′,D′ show the same overlay of markers without DAPI. A′′,B′′,C′′,D′′ show the first 2 rounds of immunolabelling of bipolar cells (PKC-β, green), glial intermediate filaments (Zrf-1, cyan), horizontal cells (CA1, purple) and Müller glia (GS, red) at different timepoints. A′′′,B′′′,C′′′,D′′′ show the last round of immunolabelling of ribbon synapses (Ribeye-A, blue), amacrine and ganglion cells (HuC/D, magenta) and Müller glia (eGFP transgene, yellow) at different, crucial timepoints of retinal development (2, 2.5, 3, and 5 dpf) showing retinal progenitors in (A-A′′′). (B-B′′′) shows nascent IPL and Müller glia formation. In (C-C′′′ and D-D′′′), IPL sublamination and Müller glia elaboration is evident. dpf: days post fertilisation, IPL: inner plexiform layer. Scale bars - 25µm for whole retina, 10µm for zoom images.

During development, nascent MG cells begin as simple unbranched radial cells at 2.5 dpf, before morphologically elaborating to a mature, highly branched structure by 5 dpf (Williams *et al*., 2010; MacDonald *et al*., 2015; Wang *et al*., 2017). MG are among the last retinal cell types to mature, integrating into neuronal circuits when neurons are undergoing robust synaptogenesis (Cepko *et al*., 1996). To determine MG specification relative to development of other retinal neurons and inner plexiform layer (IPL) formation, we used markers for MG: Zrf-1 (recognising Gfap), glutamine synthetase (GS), and the Tg(*Tp1:EGFP-CAAX*) transgenic reporter line, which labels retinal progenitors and MG (MacDonald *et al*., 2015; Kugler *et al*., 2023). We labelled amacrine cells and retinal ganglion cells with HuC/D, horizontal cells with CA-1, bipolar cells with PKC-β, and synapse formation with Ribeye-A. We observed retinal progenitors at 2 dpf (Fig. 7A-A′′′) labelled by the GFP transgene, corresponding to retinal ganglion cell (RGC) specification below the IPL, before MG genesis and onset of MG cell body basal migration at 2.5 dpf. MG are labelled by the transgene and zrf1 at 2.5 dpf, but GS labelling is not yet apparent (Fig. 7B-B′′′). At this point, the nascent IPL is present, as evidenced by the separation of the HuC/D signal and presence of ribbon synapses (Ribeye-A) (Fig. 7B′,B′′′). From this timepoint, horizontal cells are visible, marked by carbonic anhydrase (CA) below the outer plexiform layer (Fig. 7B′,C′,D′). At 3 dpf, IPL expansion and bipolar cell terminal stratification is seen (Ribeye-A and PKC-β), along with the beginning of organisation of the IPL into clear sub-laminae (Fig. 7C′-C′′′). Additionally, MG cell bodies have migrated to their final positions (Fig. 7C′′) and are marked by GS labelling. By 5 dpf, the ribbons are organised into discrete layers in the IPL, with a visible separation between the ON and OFF layers and MG have elaborated processes into this layer to provide homeostatic support functions (Fig. 7D-D′′′). As such, we used IBEX to describe retinal histogenesis, neuron migration and patterning relative to glial specification and morphogenesis across retinal development in the zebrafish. In conclusion, we were able to employ the IBEX technique and specifically designed antibody panels to describe multiple cell types at key stages of retinal development to explore their cellular relationships, which would not be possible with traditional methods.

### IBEX on frog and killifish retina tissues

To verify the versatility of the IBEX technique across various animal models, we conducted IBEX on retinas of *Xenopus laevis* tadpoles and adult African turquoise killifish. We performed IBEX on tadpole retinas to label different retinal cell types using HuC/D, PAX6, GS, GαO, cone opsin (CO), and zpr-3 antibodies (Fig. 8A, Supplementary Video 3). HuC/D labelled cells within the INL, GCL, as well as photoreceptor inner segments. PAX6 labelled cells within the GCL – either displaced amacrine cells or RGCs – while GS labelled Müller glia. GαO labelled bipolar cell bodies and densely labelled processes within the IPL. L/M cone outer segments were labelled with both CO and zpr-3. The CO antibody was raised against red opsin (lws), while zpr-3 is an anti-rhodopsin antibody which labels the green cone opsin (rhodopsin 2). IBEX was further conducted on the retinas of 8-week-old adult male African turquoise killifish (Fig. 8B, Supplementary video 4). As expected, calretinin immunoreactivity was observed in retinal ganglion cells and horizontal cells. HuC/D and PAX6 labelled amacrine and retinal ganglion cells and partial overlap of two markers was observed. Zpr-1 labelled entire double cone cells, whereas PRPH2 expression was seen in photoreceptor outer segments. Both GS and Zrf-1 (recognizing Gfap) labelled MG cells. PCNA serves as an endogenous histologic marker for the G1/S phases of the cell cycle, therefore PCNA expression serves as a marker of cell proliferation. In killifish retina, PCNA was densely expressed in the ciliary marginal zone (CMZ) and signal was also observed in the ONL. The pan-leukocyte marker Lcp-1 was observed in the INL, close to the CMZ, and within the ONL. As such, we have demonstrated that the IBEX techniques will translate to any other tissue where multiple antibodies are available.

**Figure 8.**
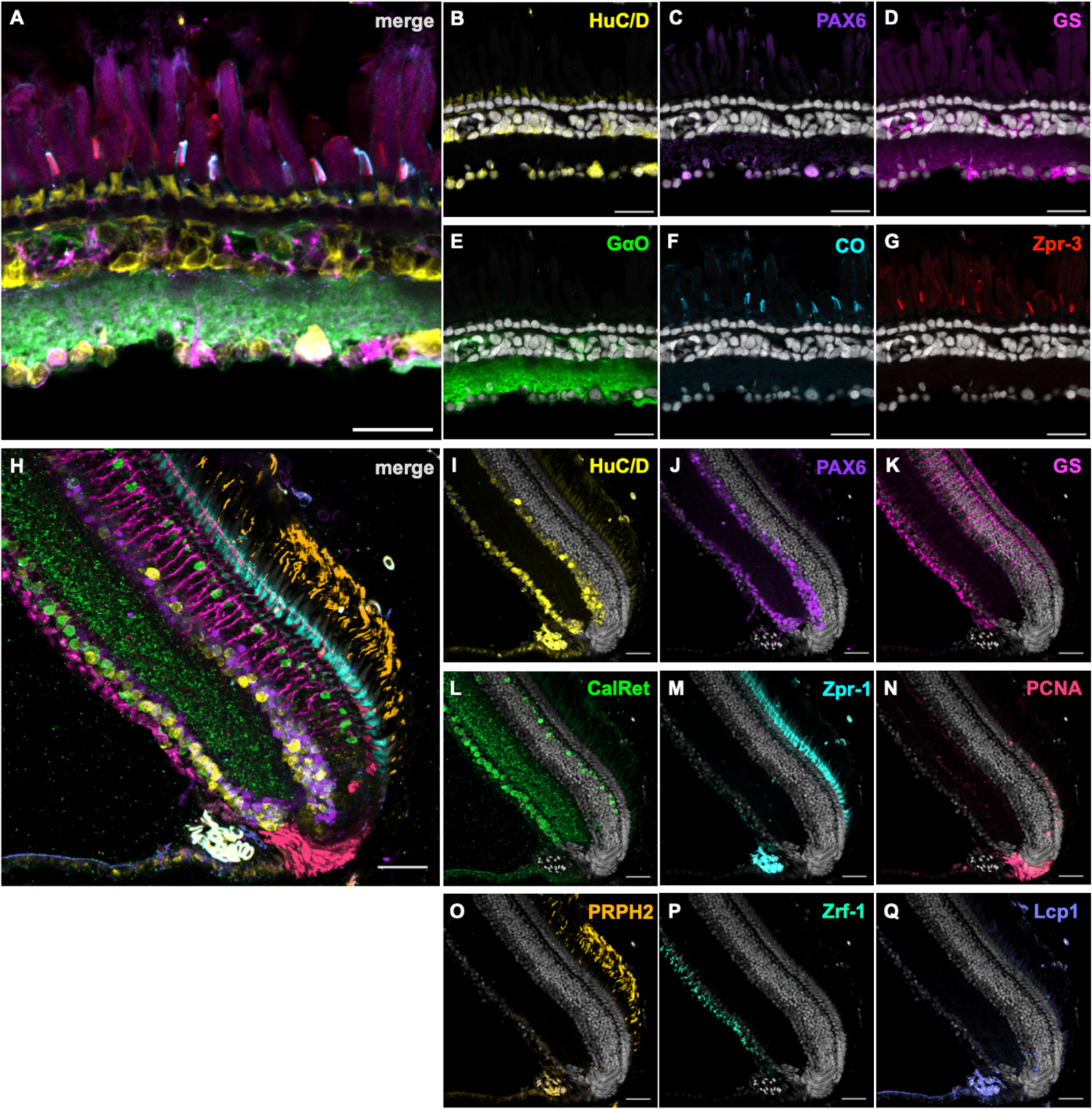
IBEX on African clawed frog and turquoise killifish retina. (A) 5-month-old *Xenopus laevis* tadpole sagittal retinal sections labelled with HuC/D (B, yellow), PAX6 (C, purple), GS (D, magenta), GαO (E, green), cone opsin (F, cyan), and zpr-3 (G, red). HuC/D labelled cells within the GCL and amacrine cell sublayer, as well as photoreceptor inner segments. PAX6 labelled primarily cells within the GCL, likely RGCs and/or displaced amacrine cells. Müller glia were labelled with GS. GαO labelled bipolar cell bodies and processes within the INL and IPL, respectively. L/M cone outer segments were co-labelled with CO and zpr-3. (H) 8-week old adult killifish retinal sections labelled with HuC/D (I, yellow), PAX6 (J, purple), GS (K, magenta), Calretinin (L, green), Zpr-1 (M, cyan), PCNA (N, red), PRPH2 (O, orange), Zrf-1 (P, light green), and Lcp1 (Q, lavender/light purple). Huc/D and PAX6 labelled cells within the GCL and amacrine cell sublayer, as seen in zebrafish and frogs. GS and zrf-1 labelled Müller glia. Calretinin labelled cells within the GCL and amacrine cell sublayer. Zpr-1 labelled entire double cone cells, and PRPH2 labelled photoreceptor outer segments. PCNA expression was observed in the CMZ and in the photoreceptor cell layer. Lcp1 was observed in the INL, close to the CMZ, and within the photoreceptor cell layer. GCL: ganglion cell layer. INL: inner nuclear layer. IPL: inner plexiform layer. RGC: retinal ganglion cell. Scale bars - 25µm for *Xenopus*; 30µm for killifish.

## DISCUSSION

There is a lack of tools available to discriminate between more than three to four cell types in a tissue simultaneously. Here, we adapted the IBEX technique to vertebrate tissues and optimised the methodology to be compatible with zebrafish cell-specific fluorescent reporter lines, wholemount IHC, and *in situ* HCR. Further, we employed IBEX to describe the relationships between glial cells and neurons and explore the entire complement of photoreceptor subtypes in the developing zebrafish retina. Additionally, we utilised IBEX to characterise the retina in killifish and *Xenopus laevis.* Therefore, IBEX is a robust method to multiplex markers and characterise cellular and molecular processes in the vertebrate model organisms.

### IBEX complements the existing zebrafish toolkit

We show that it is possible to combine IBEX with available transgenic lines with fluorescent protein expression. Zebrafish have a wealth of cell-specific transgenic reporter lines that drive transgene expression in each population of retinal cell. These have been valuable tools to characterise the development and degeneration of retinal cell types in many studies (Fadool, 2003; Bernardos and Raymond, 2006; Zolessi *et al*., 2006; Kimura, Satou and Higashijima, 2008; Vitorino *et al*., 2009; Almeida *et al*., 2014). We found that different fluorescent proteins have different bleaching success, with GFP bleaching reliably and RFP and YFP not bleaching. BFP or CFP were not assessed. Thus, it is possible to pair IBEX with transgenic lines, although this should be tested on a case-by-case basis to determine whether endogenous fluorescent proteins can be bleached, and subsequent labelling rounds adjusted accordingly. Antibodies specific for a cell type or protein of interest can be limited in non-mammalian species, and this is especially true for poorly studied or newly discovered genes. Further, studies may be interested in assessing spatial localisation of noncoding RNAs within tissues. As an alternative to antibody labelling, *in situ* hybridisation chain reaction (HCR) is a robust method to label mRNA of interest (Choi *et al*., 2010, 2016, 2018). We novelly combined *in situ* HCR with IBEX to spatially localise transcript expression for three genes with cellular markers across two cell types (three antibody labels). This can be applied to other systems to determine with high resolution which cell populations are expressing transcript for a gene of interest. Hence, combining *in situ* HCR with IBEX is a powerful technique to identify expression patterns of genes of interest by localizing gene expression to different immunolabelled cell types. WMIHC is an effective technique to study the expression pattern of proteins while preserving the 3D structure of the tissue. IBEX allows us to conduct multiple rounds of WMIHC on the same tissue. This greatly increases the amount of information we can obtain from a single tissue, is a useful extension of WMIHC to simultaneously visualise multiple proteins in their native location in tissues and explore the spatial relationships between different cell types in whole intact tissues.

### Antibody panel design and considerations

To maximise the potential of IBEX, we developed panels to label as many cell types as possible in each round of IBEX. Careful planning is required to design panels of markers (antibodies/lectins) to be used based on previous individual reactions and bleaching tests. The brightest and least efficiently bleached markers, for instance lectins or transgenes, were used in later or, ideally, last round to minimise the potential for significant leftover signal. Similarly, the weakest markers were used in early panels to increase the likelihood of strong signal detection. It is important to note that certain fluorophores are more amenable to bleaching than others, and later rounds are more susceptible to autofluorescence, particularly in the 555 channel (as in Fig. 4 and supplemental Fig. 5), where some residual inner retinal staining is observed while labelling with GNAT-2. We ensured that the fluorophores which have been validated to bleach in the original protocol (Radtke et al., 2020) were used and any new fluorophores, such as the CoraLite^®^ Plus, were tested for bleaching prior to use in IBEX (Fig. 3F). It is possible to use secondary antibodies in IBEX, and this may be required for antibodies that fail to efficiently label via micro-conjugation. However, these secondaries must be incorporated into the first round of the IBEX if recognising a species from which multiple antibodies within the designed panel are raised, as subsequent rounds using antibodies raised in that same species will lead to cross-reactivity; in the case that there is a single antibody from a specific species within the panel, the antibody can be incorporated into any round using secondary antibodies. For most fluorophores, incubation with LiBH_4_ before washing was sufficient for bleaching the fluorophore. However, in some cases it required an antigen retrieval step (i.e. heating in sodium citrate) which efficiently quenches fluorophores. Importantly, after the antigen retrieval and IBEX procedure on zebrafish retinal tissue, the nuclei appeared qualitatively similar across three rounds of IHC and imaging (Fig. S2) and were easily aligned using the SimpleITK registration software.

### Potential of IBEX and multiplexing in other species

IBEX can also be applied to any species where multiple cell-specific antibodies, fluorescent reporter transgenes, and/or fluorescent *in situ* hybridisation techniques are available. However, these techniques and antibody panels will need to be validated on a case-by-case basis. These methodologies, some developed here, will be especially valuable where multiple different markers are required to confirm the identity of a cell, such as immunological studies. We also demonstrate the benefits of IBEX to characterise multiple cell types in unconventional model organisms, such as the killifish and *Xenopus laevis,* in which there are few transgenic reporter lines. IBEX allows us to maximise the information that can be obtained from a single piece of tissue, a huge benefit for study of rare species or tissue that is difficult to access. Furthermore, it can be used to potentially identify subpopulations that would not be possible with conventional IHC in such organisms. Labelling multiple markers on the same tissue will also have important implications for animal ethics and 3Rs (Replacement, Reduction, and Refinement) initiatives as the number of animals required for statistically significant phenotypic data is greatly reduced. Performing multiple rounds of IHC on the same tissue produces rich datasets where the interactions between multiple cell types can be explored. As such, multiplexing techniques are not only powerful for exploring cellular relationships but also critical for efforts to reduce animal numbers in scientific research.

### Limitations and future work

There were some limitations to the IBEX approach. Micro-conjugations can be unreliable on occasion (i.e. not all antibodies successfully conjugate), there is a need for a relatively large volume of validated primary antibody (although much less than when performing a primary conjugation), and the technique works most effectively when the protein in question is highly abundant. Hence, when used for proteins that have low expression or low affinity for antibodies, the concentrations may have to be increased over traditional methods. Many antibodies do not have their exact antigen identified or validated in non-mammalian systems, such as zebrafish. However, as we were only looking at broad cell type distributions, rather than targeted molecular events, validation of each antigen was not necessary for this study. When possible, we did utilise antibodies with known/validated antigens. When using unvalidated antibodies, we selected those that had conserved labelling patterns and included other methods – such as transgenic lines and *in situ* hybridisation – to complement the labelling and provide information about specificity. Transgenic lines with RFP/YFP that do not bleach (such as the Tg(*rho:YFP*) line used for the developmental series in Fig. 7) can still be used if required but would reduce the number of channels available for successive imaging rounds. Finally, there are likely limitations on the number of immunolabelling rounds that can be performed without tissue damage or visible residual signal from previous rounds. However, we have conducted four rounds of IBEX on retinal cryosections and there has been no noticeable degradation of tissue integrity or signal loss. As such, it may be possible to >20 markers on a single zebrafish cryosectioned tissue, as there have been 20 rounds and 66 antibodies reported in human lymph nodes (Radtke *et al*., 2020).

The IBEX technique is compatible with standard confocal as well as epifluorescence microscopy. Here, we principally used a standard confocal microscope with four excitation laser wavelengths (405, 488, 555, and 647) that allowed a maximum of three antibodies plus DAPI in each panel. However, this can be expanded upon if your microscope has additional spectral capabilities (e.g. tuneable or white light laser) and antibodies are visualised with additional fluorophores via conjugation or secondary antibodies. To this end, we also conducted IBEX using an epifluorescence microscope with Z-stack capabilities and expanded spectral excitation properties to visualise 10 markers on the same tissue (Fig. S3). As such, this technique will be applicable to any fluorescence microscopes available where multiple channels can be acquired on the same tissue.

In conclusion, this technique can be a powerful method to explore cellular relationships and biological processes in complex multicellular tissues in the context of development, ageing or disease.

## MATERIALS AND METHODS

### Animals

Adult fish were housed in the animal facility at the University College London on a 14:10 hour light/dark cycle at 28°C, following previously established husbandry protocols (Westerfield, 1993). Experimental procedures were conducted in accordance with the UK Home Office Animals (Scientific Procedures) Act 1986 (zebrafish project license PPL: PP2133797, held by R.B.M, and killifish PPL: PP7179495 held by Dr Elspeth Payne). Zebrafish embryos were obtained by light-induced spawning, collected in E3 buffer (5 mM NaCl, 0.17 mM KCl, 0.33 mM CaCl2, 0.33 mM MgSO4) with/without methylene blue, and maintained in an incubator at 28.5°C till use.

*Xenopus laevis* (African clawed frog) use was approved by the University of Alberta Animal Use and Care Committee (AUP00004203) and carried out in accordance with the Canadian Council on Animal Care. Frogs were housed at 18°C under a 12-hour cyclic light schedule (7:00-19:00; 900-1200 lux).

### Animal strains

Wildtype zebrafish embryos (ABTL/Tübingen) were used for the adaptation of the IBEX technique on section IHC, WMIHC and combined HCR/IHC. Tg(*GFAP:GFP*) (Bernardos and Raymond,2006), Tg(*vsx1:GFP*)^nns5^ (Kimura, Satou and Higashijima, 2008), Tg(*TP1:Venus-Pest*) (Ninov, Borius and Stainier, 2012), Tg(*tp1bglob:eGFP-CAAX*) (Kugler *et al*., 2023) and Tg(*ptf1a:dsRed*)^ia6^ (Jusuf *et al*., 2012) were used to optimise the protocol for IBEX using transgenics. Tg(*rho:YFP*)^gm500^ (White *et al*., 2017), Tg(*tp1bglob:eGFP-CAAX*) embryos were used at different timepoints to study development of neurons and glia. Wild type killifish adults (GRZ) and wild type *Xenopus laevis* were used for adaptation of IBEX on retinal sections.

### Preparation of retinal sections

Animals at the desired stages were overdosed with 0.4% Tricaine and fixed in 4% paraformaldehyde overnight at 4°C. Tissues were washed 3 times for 5 min in PBS and then immersed in either 20% (frogs) or 30% (fishes) sucrose in PBS and allowed to sink overnight. Samples were embedded in OCT (Sigma Aldrich, Cat. No. SHH0024) and frozen at −80°C. Frog eyes were obtained by B.J.C. Froglets aged 145 days post-fertilization were euthanised by overdose with tricaine (0.5% until unresponsive to toe pinch), decapitation, and then pithing. Whole eyes were fixed in 4% PFA + 3% sucrose overnight, cryoprotected in 20% sucrose overnight with gentle shaking, and then shipped to UCL in 20% sucrose for further processing. SuperFrost™ Plus Adhesion Microscope Slides (Epredia, Cat. No. J1800AMNZ) were coated evenly with 5μL chrome alum gelatin (Newcomer Supply, Part# 1033C) and dried in an incubator at 60°C for one hour, to minimise loss of tissue over multiple rounds of immunolabelling. Retinal sections of embedded tissues were sectioned onto these slides at a thickness of 12-14 µm using a cryostat (Leica CM1950) and left at room temperature (RT) to dry overnight. Slides were then stored at −80°C.

### Antibody micro-conjugation

Each antibody (Table 1) was tested with the FlexAble micro-conjugation kits at the recommended concentration (0.5 µg) per slide. However, this did not label cells efficiently in retinal sections. Doubling the concentration of primary antibody was found to stain tissue more effectively. 1 µg of each primary antibody was combined with 2 µL of FlexAble linker protein for the desired fluorophore, and the volume was made up to 16 µL with the provided buffer, following kit recommendations. This solution was incubated for 5 minutes in the dark at RT, and then 4 µL of quencher was added and left to incubate in the dark at RT for a further five minutes, according to the manufacturer’s protocol. The entire reaction volume for each antibody was used for subsequent steps.

**Table 1.**
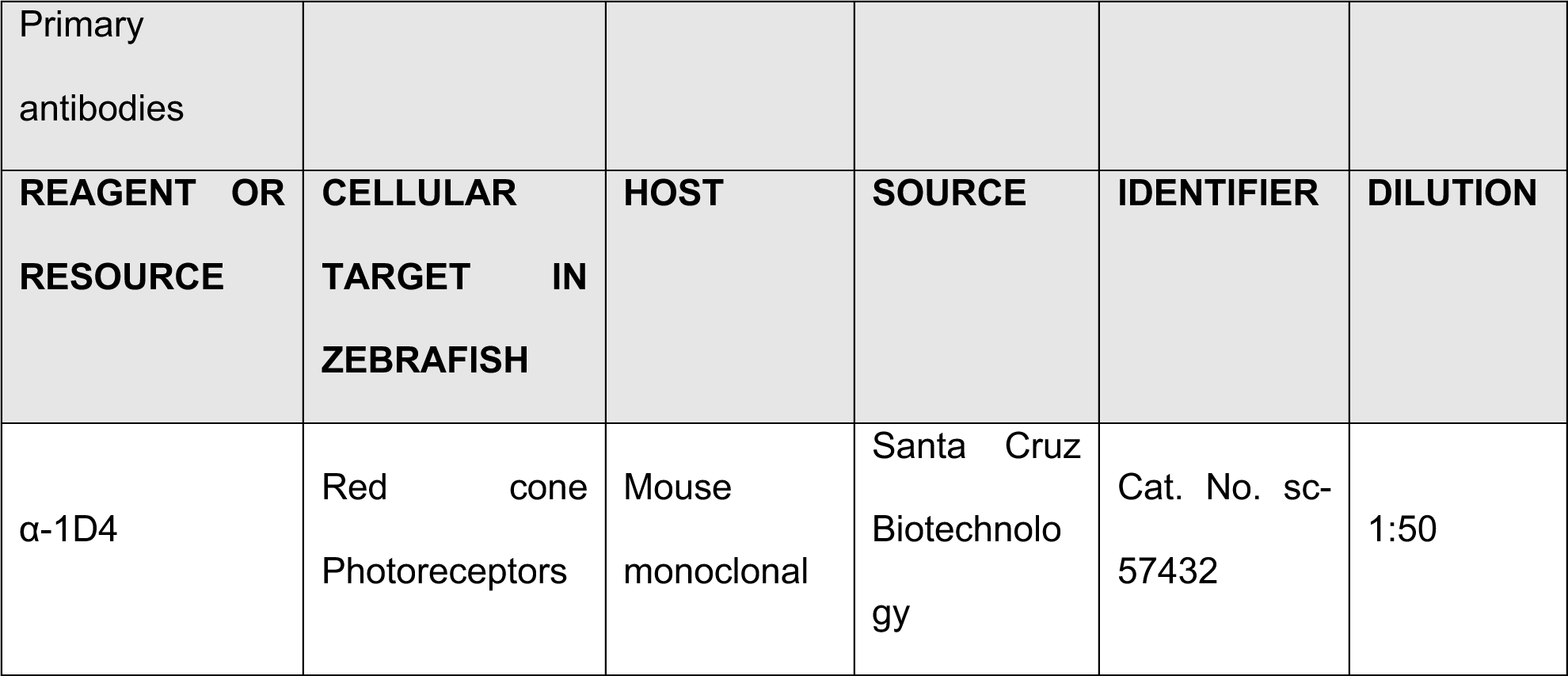
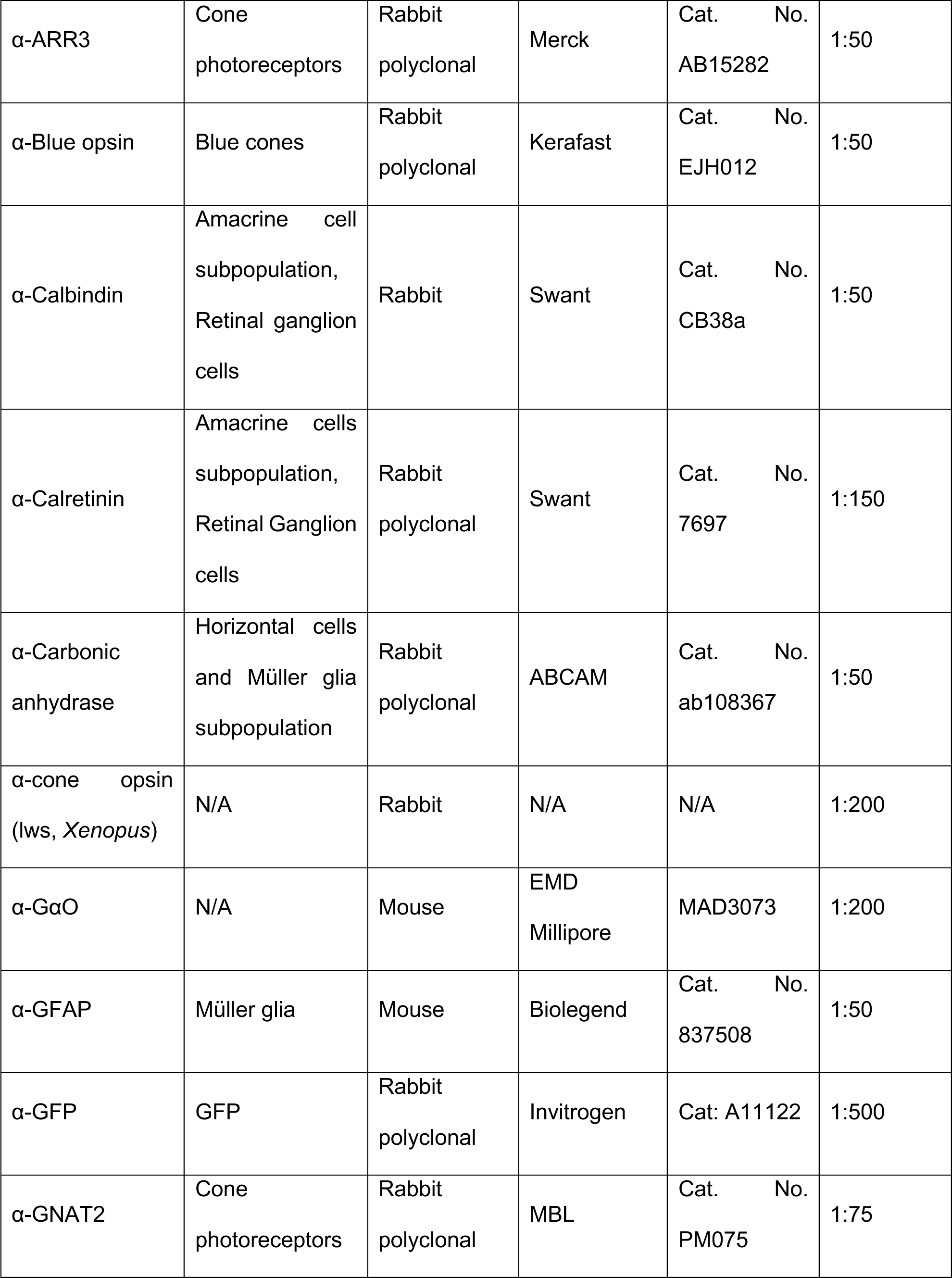
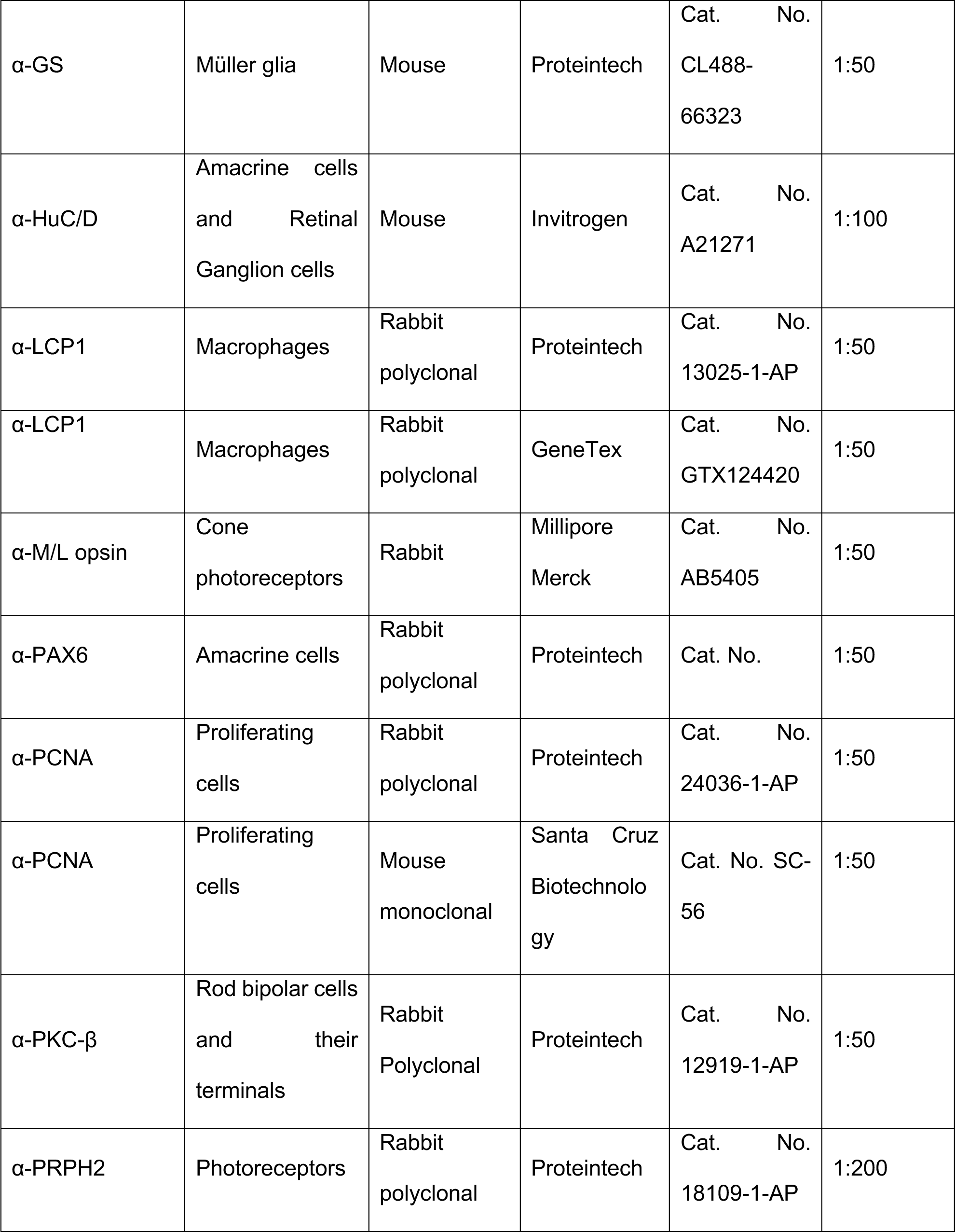
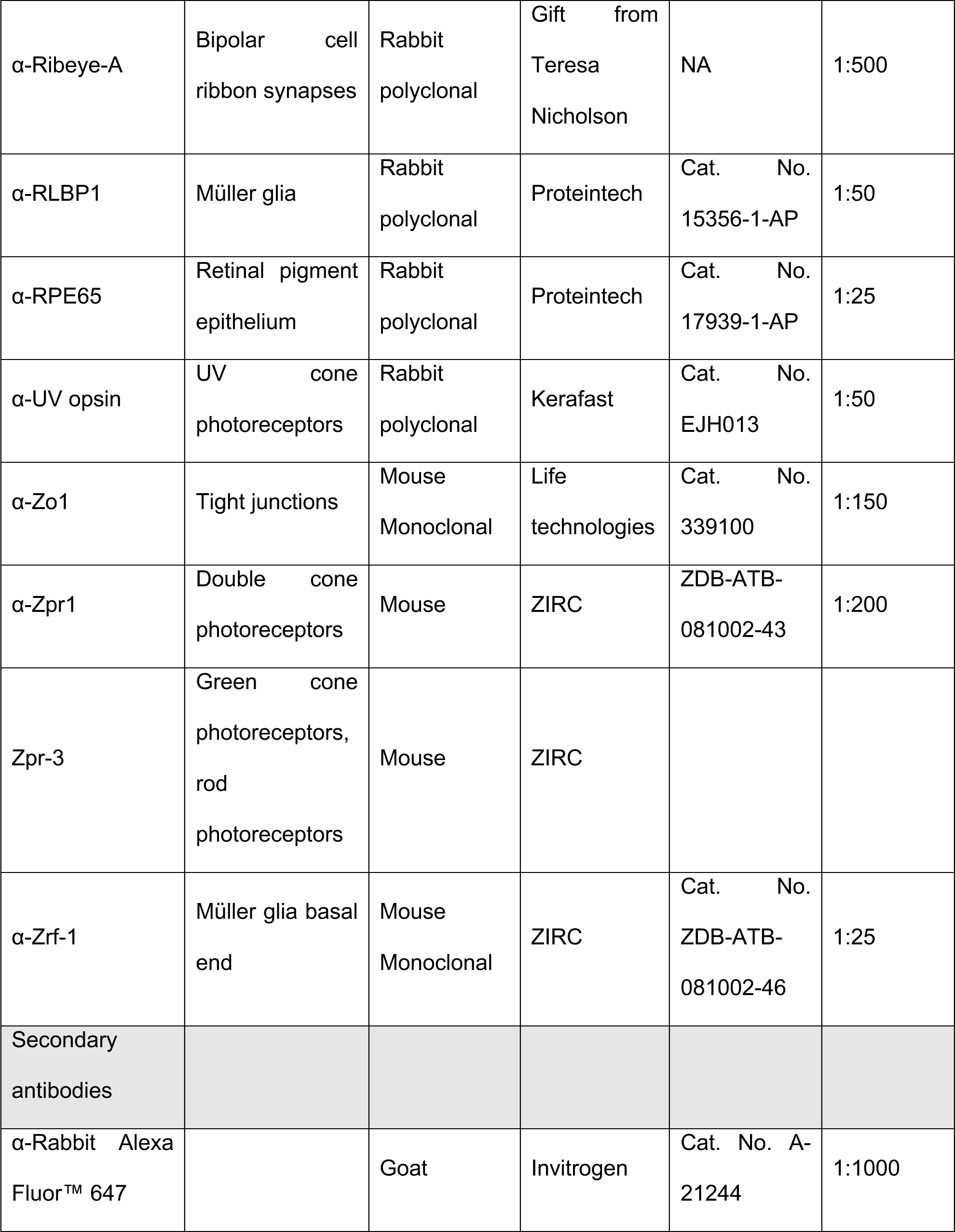
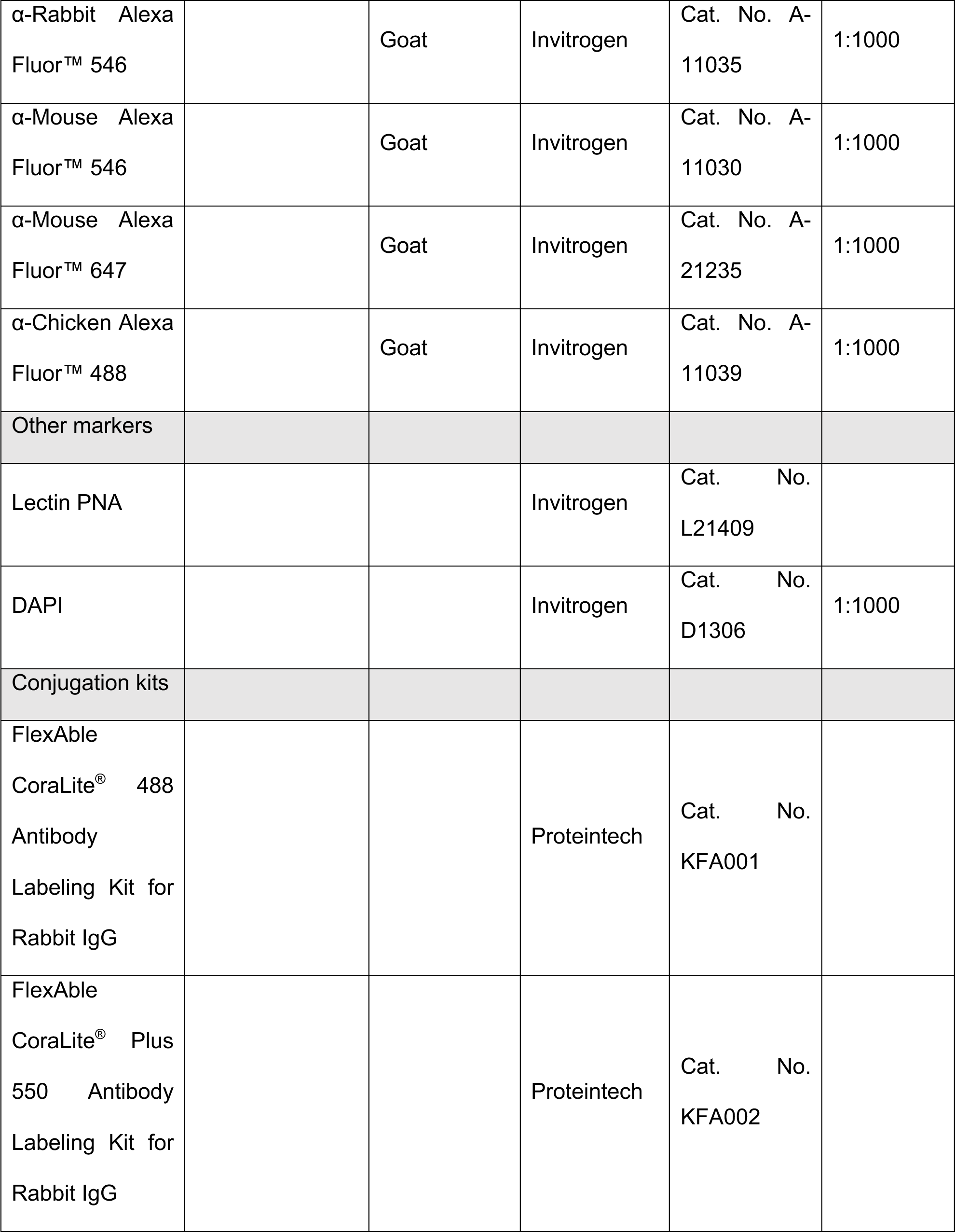
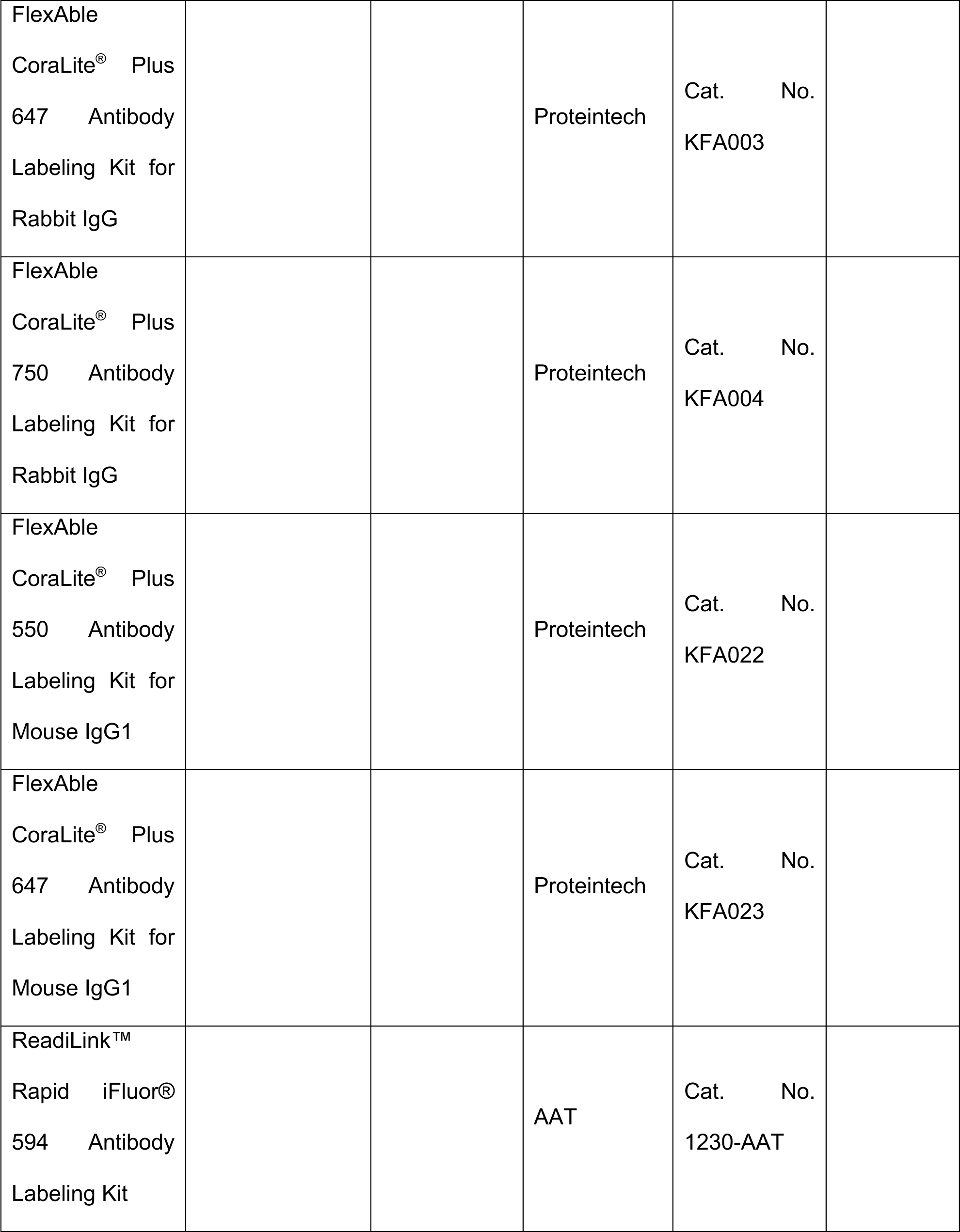
Antibodies used for immunohistochemistry and IBEX.

### IBEX technique

This method is an adapted version of the original IBEX protocol (Radtke *et al*., 2020) and an overview is given in Fig. 2. The sections were rehydrated in PBS for 5 minutes at RT. Antigen retrieval was then performed by heating the slides for 20 minutes in 10 mM sodium citrate (pH 6). This step quenched the signal in most of the zebrafish GFP transgenic lines tested, but the GFP can be boosted in later rounds of immunolabelling using a primary antibody against GFP. The sections were blocked for 1 hour in block solution (10% goat serum, 1% BSA, 0.8% Triton X, 0.1% Tween, made up with PBS) at RT. Primary antibodies were conjugated to fluorophores as described above. The slides were incubated with of the first round of antibodies, diluted appropriately in block solution, at 4°C overnight. Following three 20-minute washes with PBS, secondary antibodies were added, if needed. Slides were then incubated at RT for two hours or overnight in 4°C and washed after with PBS three times for 10 minutes. Slides were mounted in Fluoromount G mounting media (Cat. No. 00-4958-02, Invitrogen) and imaged on a Leica THUNDER imager, Leica SP8 confocal microscope, or Zeiss LSM 900 inverted confocal using 4-5 channels: 405, 488, 550, 647 and 750 nm.

After image acquisition, slides were placed in a 50 mL falcon tube filled with PBS and left until the coverslip fell off, and then washed three times to remove the mounting media. Fluorophores were quenched by incubating slides in 150 µL of lithium borohydride (LiBH_4_, 16949-15-8, STREM) solution (1-2 mg/mL) under direct white light (minimum of 300 lux). The slide was then washed three times for 10 minutes in PBS before the next round of antibodies was added and steps were repeated as above.

### IBEX for combined *in situ* HCR/IHC on sections

HCR probes for *cyp26a1* and *vsx1* were kindly gifted by Takeshi Yoshimatsu, while the probe set for *glula* was designed using a custom script (Trivedi and Powell, unpublished) and ordered from Life Technologies, ThermoFisher. HCR amplifiers (Alexa Fluor 488, Alexa Fluor 546, and Alexa Fluor 647), and buffers (hybridisation, wash, and amplification) were purchased from Molecular Instruments (https://www.molecularinstruments.com/). A published *in situ* HCR protocol (Choi *et al*., 2018) was adapted for retinal zebrafish sections. Slides were rehydrated in PBS for 5 minutes and then treated with 250 µL proteinase K (20µg/mL) for 10 minutes at RT. They were washed twice with PBS + 0.1% Tween (PBST) and post-fixed with 250 µL of 4% paraformaldehyde for 20 minutes at RT. Slides were washed 3 times for 5 minutes with PBST and incubated in 150 µL of probe hybridisation buffer at 37°C for 30 minutes. Sections were then incubated overnight at 37°C in probe solution (4 µL of each probe set, made up to 150 µL in probe hybridisation buffer). Excess probes were removed by washing slides 4 times for 15 minutes in probe wash buffer at 37°C, and then 2 times for 5 minutes with 5X SSCT buffer at RT. Slides were pre-amplified in 150 µL of amplification buffer for 30 minutes at RT. 4 µL each of hairpin 1 and hairpin 2 amplifier were snap cooled by heating to 95°C for 90 seconds and cooling to RT. These were then added to the slides in 150 µL of amplification buffer and incubated overnight at RT in the dark. Slides were washed 4 times for 5 minutes in 5X SSCT buffer and mounted in Fluoromount G media. Sections were imaged on the Zeiss LSM 900 with a 40x immersion oil objective (Na 1.1) using 4 channels: 405, 488, 546, and 647 nm. After image acquisition, slides were heated to 60°C in sodium citrate (pH 6) for 20 minutes to quench fluorophores. IHC was then repeated on sections as described above.

### IBEX for WMIHC

Wildtype zebrafish embryos were treated with 0.0045% phenylthiourea from 6 hpf to prevent pigment formation and at 5 dpf, were overdosed with 0.4% Tricaine and fixed in 4% paraformaldehyde overnight at 4°C. They were washed in PBST and heated in 10mM sodium citrate (pH 6) at 70°C for 15 minutes. Samples were washed twice for 10 minutes in PBST, twice for 5 minutes in distilled water and incubated with ice cold acetone for 20 minutes at −20°C. This was followed by three 5 min washes in PBS and incubation in blocking solution (10% goat serum, 0.8% Triton X-100, 1% BSA in PBST) for 2 hrs at RT. Micro-conjugation of the antibodies was carried out as described above, doubling the volume of antibody used for sections. Embryos were incubated in antibody solution with DAPI, diluted with blocking solution, at RT overnight with gentle agitation at room temperature. After three 1-hour washes in PBS + 1% Tween, embryos were mounted in molten 1% low melting point agarose in PBS, in a glass bottomed dish. Once hardened, they were covered with 1X PBS and imaged on the Zeiss LSM 900 with a 40x immersion oil objective (Na 1.1) using 4 channels: 405, 488, 546, and 647 nm. Embryos were then extracted from the agarose and bleached with LiBH_4_ solution mentioned above in individual tubes for 2 hours at room temperature. The antibody incubation process was repeated, keeping embryos separate in order to overlay correct images during registration. They were then imaged as mentioned above, keeping imaging parameters like stack size and step size consistent between rounds.

### Optimization of quenching of fluorophores

Fluorophores react differently to bleaching with LiBH_4_, and therefore, we used several different methods to bleach signal between rounds of labelling. CoraLite micro-conjugated antibodies bleached after 30 minutes of LiBH_4_ treatment under bright light. The same antibodies required 2 hours of incubation in LiBH_4_ solution to observe reduction in signal in WM IHC. In cases where Alexa Fluor 555 secondary antibody was used, 30 minutes of heating at 60°C in sodium citrate solution (pH 6) was needed to quench the fluorescence. For transgenic lines, antigen retrieval by boiling slides in sodium citrate solution (pH 6) for 20 minutes quenched the fluorescent protein, but when boosted with the anti-GFP primary and Alexa Fluor secondary antibodies, the signal was not reduced even after antigen retrieval. Hence, fluorophores must be individually tested to assess suitability for use with IBEX, and ones that do not show reduction in signal used in the last round of immunolabelling.

### Image processing and alignment

Imaging parameters such as stack size, number of steps, step size, scan speed, and resolution were kept consistent across all rounds of imaging. Once all rounds were completed, Imaris (Oxford Instruments) was used to process the images. Brightness, contrast, and colours were adjusted and filters, such as Gaussian, were applied where appropriate. Once all images were processed, the SITK-IBEX registration code (Radtke *et al*., 2020) was used to register them, using DAPI as the alignment channel. Maximum projection images were obtain using the snapshot feature while in 3D viewer or by using the orthogonal slicer tool.

## ACKNOWLEDGEMENTS

We would like to thank Dr. Mariya Moosajee for the kind gift of *Tg(rho:YFP)^gm500^* embryos, and Dr. Chintan Trivedi for allowing use of his custom HCR probe design script. We also thank Dr. Elspeth Payne for providing access to killifish tissue and UCL BRU staff for care and maintenance of animals used in this study. This study was supported by BrightFocus Macular Degeneration Postdoctoral Fellowships to N.C.L.N. (M2022002F) and B.J.C. (M2021001F), University of Alberta startup funds to B.J.C, a Wellcome Trust Clinical Research Career Development Fellowship (224586/Z/21/Z) and Moorfields Eye Charity equipment grant to C.J.C and a BBSRC David Phillips Fellowship (BB/S010386/1) and a Moorfields Eye Charity PhD Studentship (GR001503) to R.B.M.

## SUPPLEMENTAL FIGURES

**Supplemental Figure 1.**
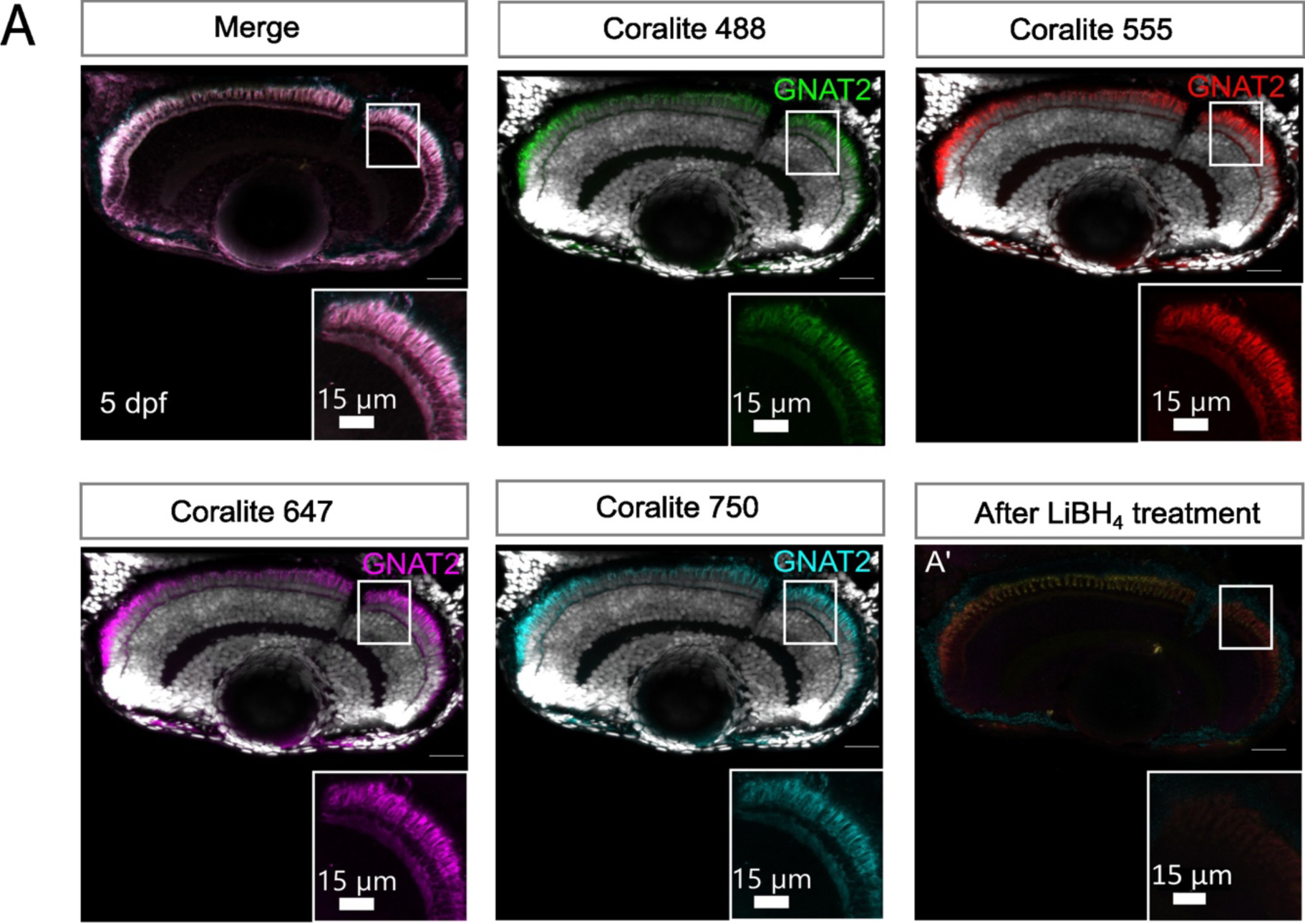
(A) Epifluorescence images of cone photoreceptor cells immunolabelled with the GNAT2 antibody on the same single retinal section. Rabbit α-GNAT2 is conjugated to Coralite 488, Coralite 550, Coralite 647 and Coralite 750. (A′) Decreased signal after bleaching the sample with LiBH4 under bright light. Scale bars - 40µm for whole retina, 15µm for zoom images.

**Supplemental Figure 2.**
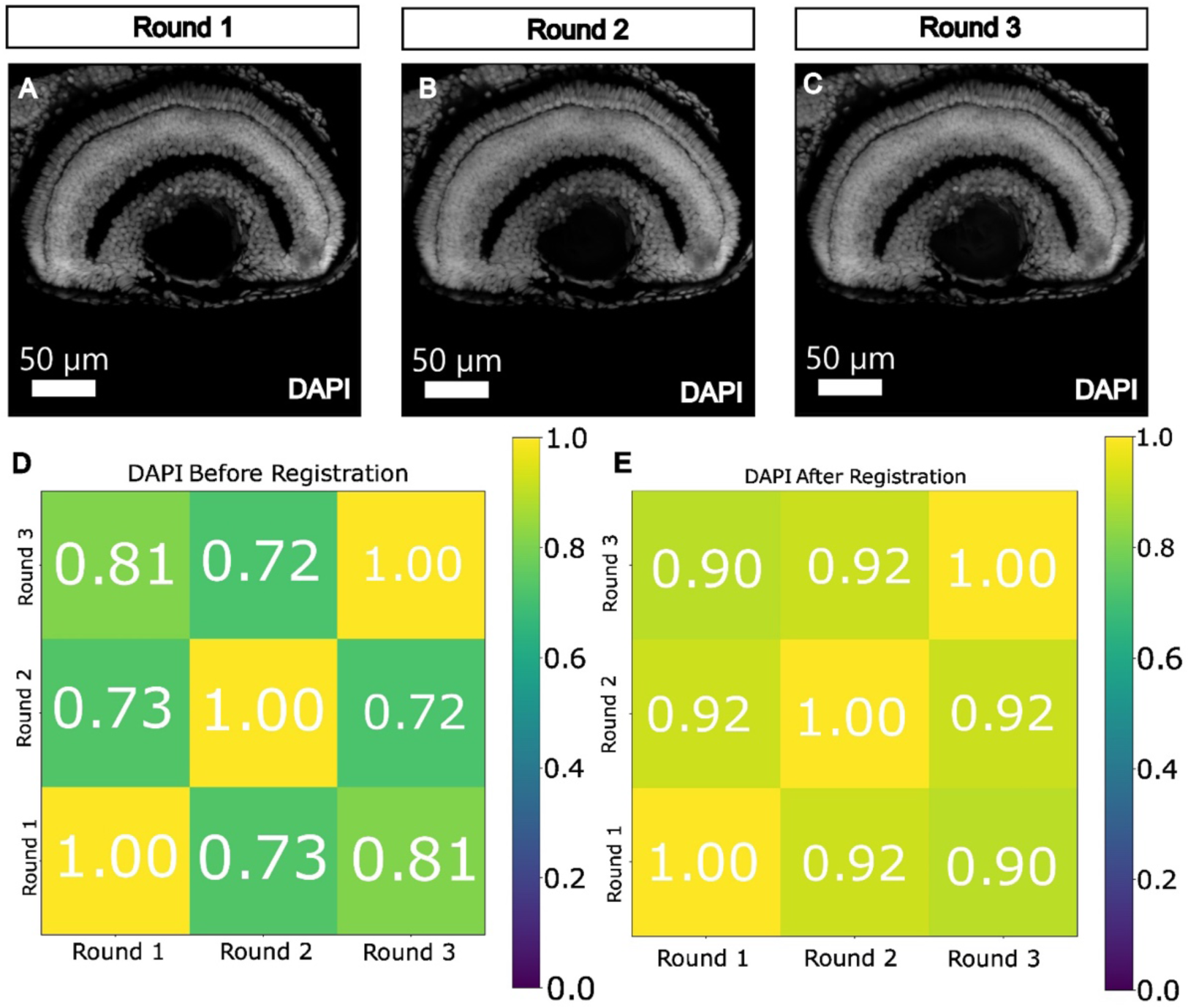
Three round IBEX run on 5 dpf zebrafish retina cryosections with Z-stacks using a 63x objective and 2x optical zoom on the Leica SP8 confocal microscope (three separate technical repeats were performed). (A-C) Maximum projections of DAPI staining during the imaging of rounds 1-3, respectively. (D) Correlation matrix for DAPI staining across the rounds before Simple ITK registration of Z-stacks. (E) Correlation in DAPI staining across the rounds after affine registration of Z-stacks. Scale bars - 50µm.

**Supplemental Figure 3.**
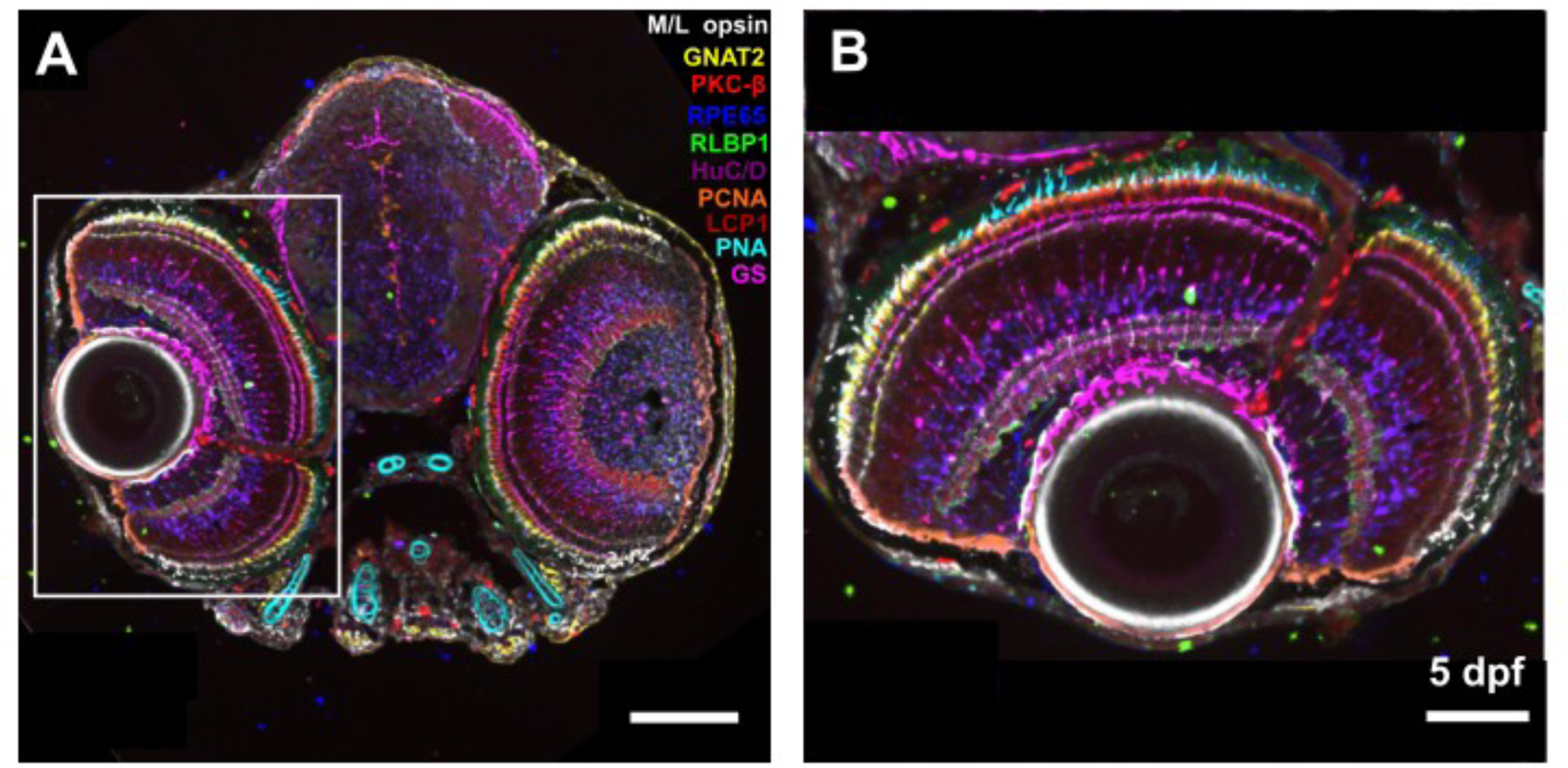
Three round IBEX run using the Leica THUNDER imager with a 40x air objective following instant computational clearing.(A) Epifluorescence images of 5dpf zebrafish retinal section immunolabelled with a panel of 9 antibodies and a lectin stain: RPE65 (dark blue), GS (magenta), PKC-β (red), HuC/D (purple), PCNA (orange), GNAT2 (yellow), RLBP1 (green), Lcp-1 (maroon), M/L opsin (white) and PNA stain (cyan). (B) Zoom in of region of interest indicated in A, with merge of the 10 antibodies used. Scale bar - 80µm (A) and 40µm (B).

**Supplemental Figure 4.**
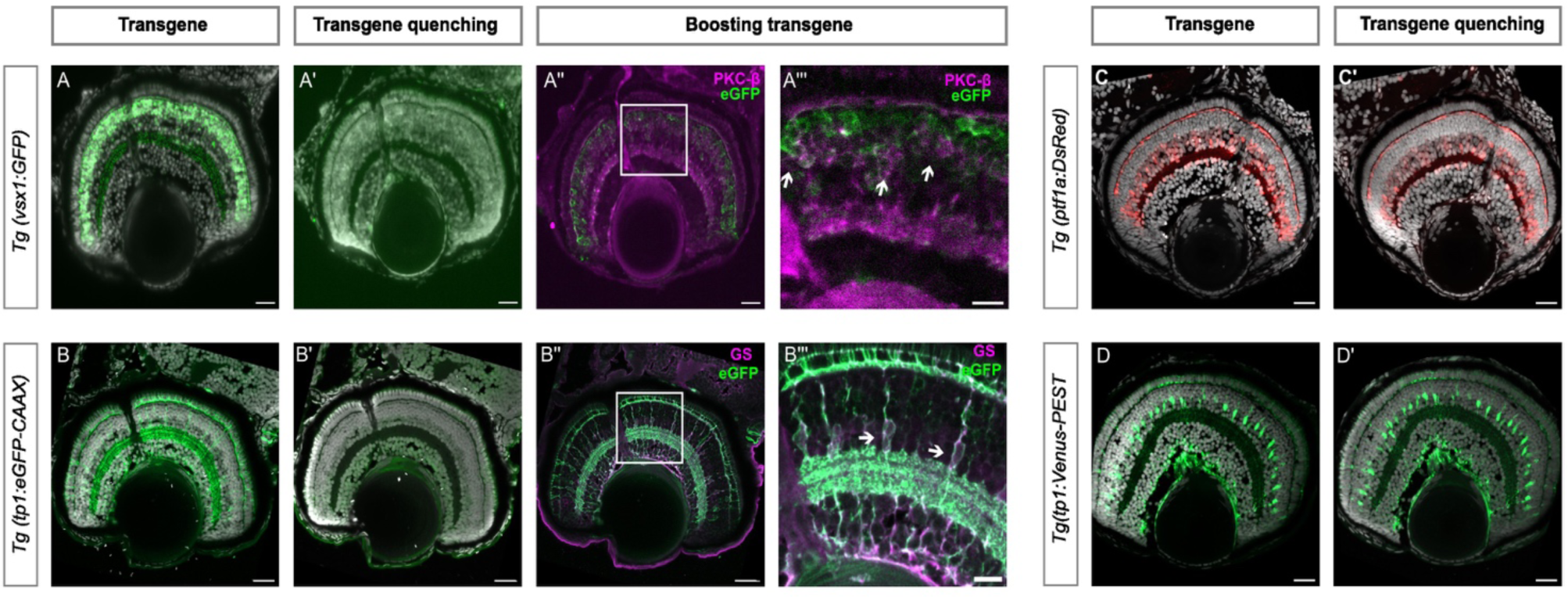
IBEX is compatible with transgenic fluorescent reporter lines. (A) Images of 5 dpf cytosolic GFP (Tg(*vsx1:GFP*) transgenic labelling bipolar cells) positive retinal sections without any treatment. (A′) Images of the same section after sodium citrate treatment, showing reduced signal. (A′′) Section imaged after boosting transgene with anti-GFP antibody and co-labelling with PKCβ to confirm specificity of boosted transgene. (A′′′) Zoom in of region of interest in (A′′). (B) Confocal images of 5 dpf membrane bound GFP (Tg(*tp1:eGFP-CAAX*) transgenic labelling MG membranes) positive retinal sections without any treatment. (B′) Images of the same section after sodium citrate treatment, showing reduced signal. (B′′) Section imaged after boosting transgene with anti-GFP antibody and co-labelling with GS to confirm specificity of transgene (zoom: arrows). (B′′′) Zoom of region of interest indicated in (B′′). (C, D) Confocal images of RFP (Tg(*ptf1a:Dsred*) transgenic) and YFP (Tg(*tp1:Venus-PEST*) transgenic retinal sections without treatment. (C′,D′) RFP and YFP transgenes do not show any change in signal after sodium citrate treatment. dpf: days post fertilisation, MG: Müller glia. Scale bar - 25µm.

**Supplemental Figure 5.**
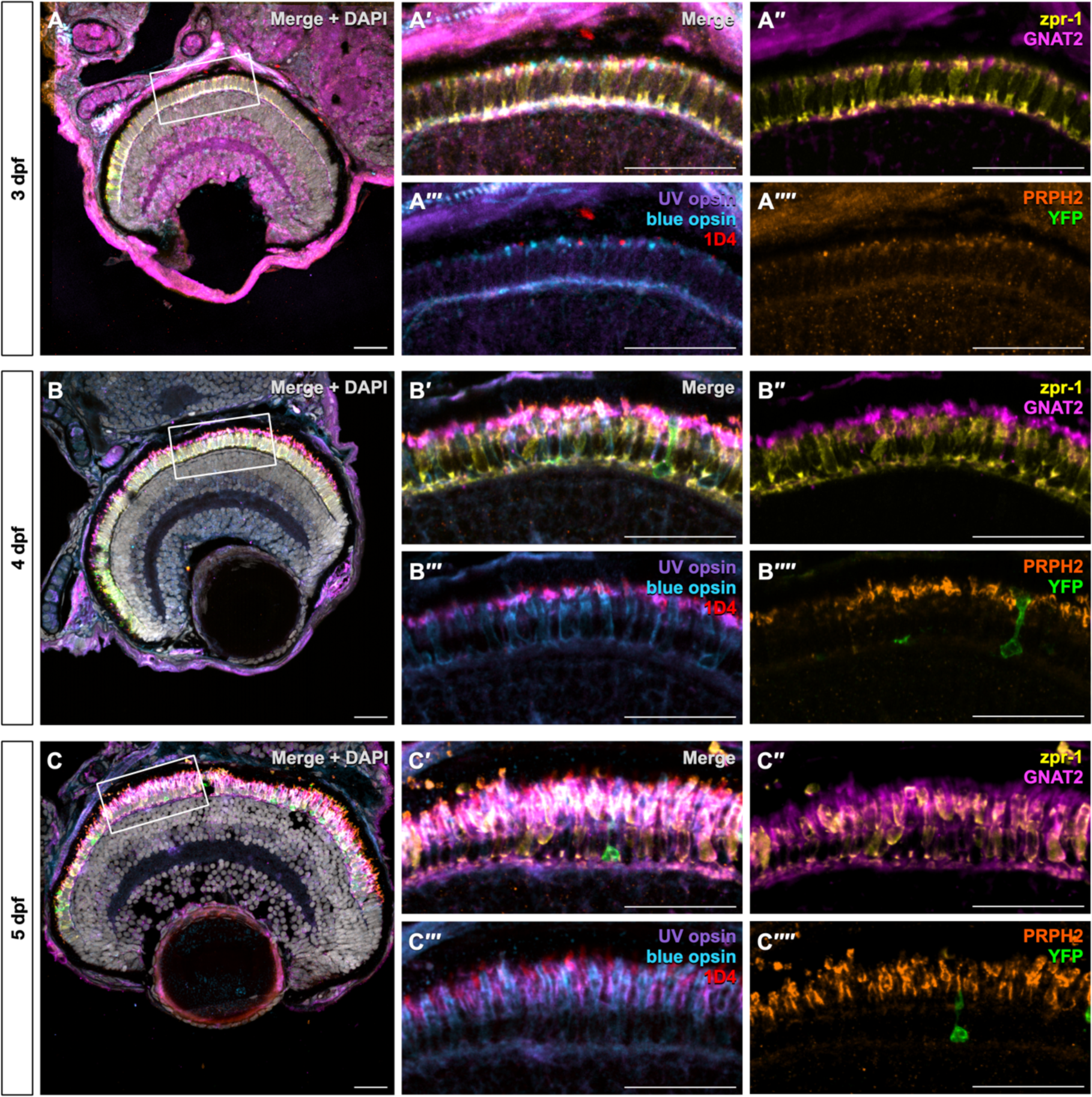
Visualisation of all photoreceptor subtypes during zebrafish development using IBEX. Sagittal sections of Tg(*rho:YFP*) embryos labelled with zpr-1 (yellow), GNAT2 (pink), UV opsin (purple), blue opsin (blue), 1D4 (red), and PRPH2 (orange) at 3 (A), 4 (B), and 5 (C) dpf. YFP is shown in green. (A′, B′, C′) Show zooms of photoreceptors will all labels merged, without DAPI; (A′′, B′′, C′′) shows zpr-1 and GNAT2 labelling; (A′′′, B′′′, C′′′) shows UV, blue, and red opsin (1D4) labelling; and (A′′′′, B′′′′, C′′′′) shows PRPH2 labelling and YFP. (A) 3 dpf retinas have small, newly developing outer segments visible by UV opsin, blue opsin, red opsin (1D4), and PRPH2 labelling. Entire cone cell bodies are visible by GNAT2 labelling, while red and green cone cell bodies are visible by arrestin 3a (zpr-1) labelling. Newly formed rods are visible at the periphery of the retina by YFP labelling. (B) 4 dpf embryos have longer outer segments with increased PRPH2 staining indicative of disc structure. (C) 5 dpf embryos have long outer segments that are visibly starting to taper into more of a “cone-like” morphology. Scale bars - 25µm.

**Supplemental Figure 6.**
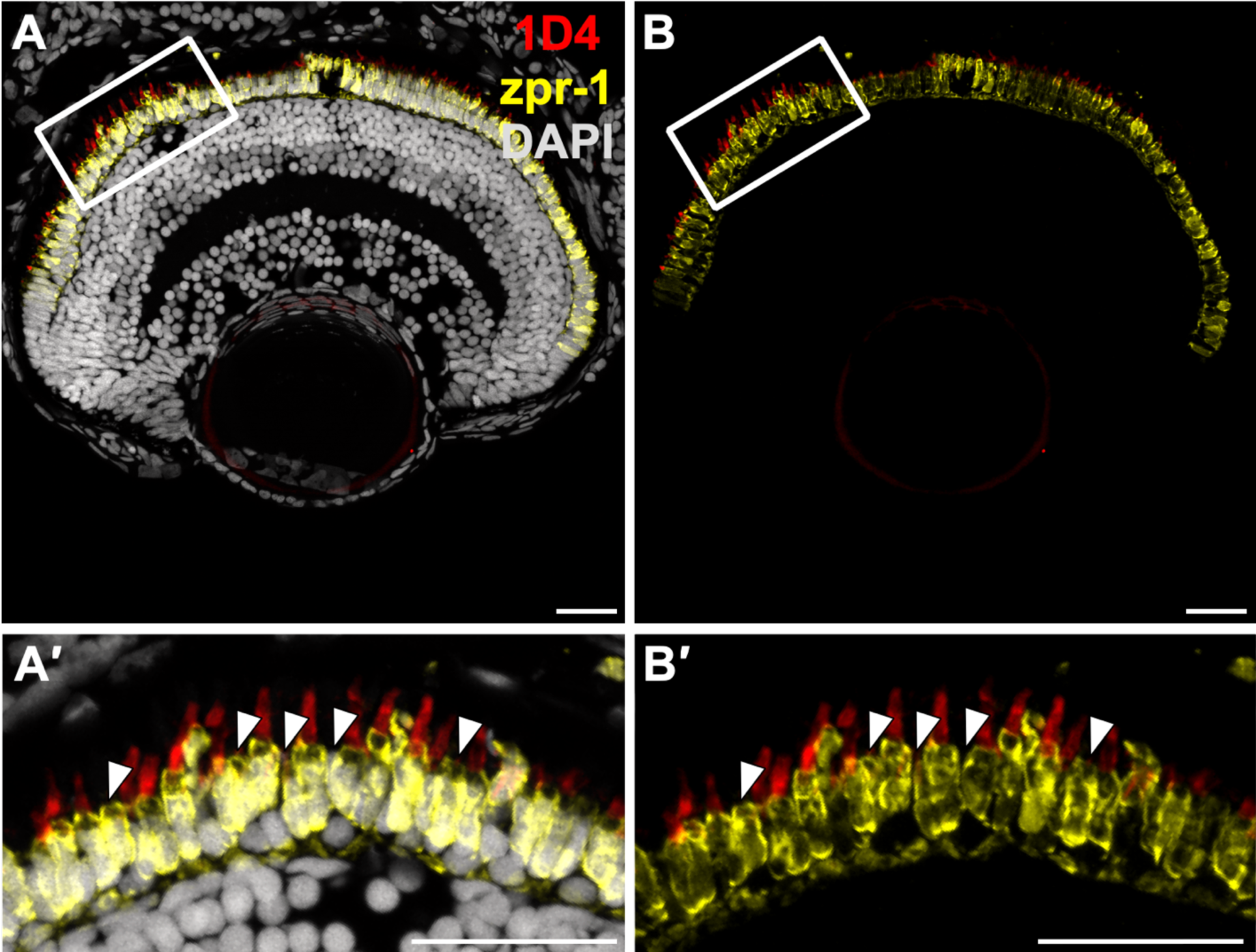
Discrimination of green cones in 5 dpf zebrafish retina. Red cones are positive for both zpr-1 and 1D4 (red opsin), whereas green cones are positive for only zpr-1 (arrowheads). Scale bars - 25µm.

